# Complete sequences of the velvet worm slime proteins reveal that slime formation is enabled by disulfide bonds and intrinsically disordered regions

**DOI:** 10.1101/2022.02.18.480993

**Authors:** Yang Lu, Bhargy Sharma, Wei Long Soon, Xiangyan Shi, Tianyun Zhao, Yan Ting Lim, Radoslaw M. Sobota, Shawn Hoon, Giovanni Pilloni, Adam Usadi, Konstantin Pervushin, Ali Miserez

**Author notes:** Y.L and B.S contributed equally to the manuscript.

## Abstract

The slime of velvet worms (Onychophora) is a strong and fully biodegradable protein material, which upon ejection undergoes a fast liquid-to-solid transition to ensnare prey. However, the molecular mechanisms of slime self-assembly are still not well understood, notably because the primary structures of slime proteins are yet unknown. Combining transcriptomic and proteomic studies, we have obtained the complete primary sequences of slime proteins and identified key features for slime self-assembly. The high molecular weight slime proteins contain Cys residues at the N- and C-termini that mediate the formation of multi-protein complexes via disulfide bonding. Low complexity domains in the N-termini were also identified and their propensity for liquid-liquid phase separation established, which may play a central role for slime biofabrication. Using solid-state nuclear magnetic resonance, rigid and flexible domains of the slime proteins were mapped to specific peptide domains. The complete sequencing of major slime proteins is an important step towards sustainable fabrication of polymers inspired by the velvet worm slime.

## Introduction

Many living organisms secrete structural fibrous materials in their immediate external environment to secure locomotion^1^, attach themselves to solid substrates^2^, or capture prey^3^. These abilities provide valuable lessons for biomimetic and sustainable biopolymer processing^4^. A classic example is spider silk, whose bio-fabrication of mechanically-tough fibers has been investigated for several decades^5^. An organism that has more recently garnered interest as a model system for biopolymer fabrication is the velvet worm (Onychophora).

Onychophora are carnivorous invertebrates that are placed in their own phylum and have existed for more than 500 million years. The preying mechanism of velvet worms is unique, consisting in rapid secretion of a highly adhesive slime out of oral papillae to capture small insects (**Fig. 1**). The secreted slime has many remarkable characteristics: *(i)* it exhibits high extensibility and tensile strength (ultimate tensile strength of 101.9 ± 20.1 MPa^6^); *(ii)* it rapidly phase-separates from a concentrated dope solution into the adhesive slime as soon as it is ejected out of the slime gland; *(iii)* it is water soluble and *(iv)* it can reversibly form fibers after being re-dissolved. The slime is mainly composed of highly hydrated proteins (90% water in the wet slime)^7^, with a small fraction of lipids. Separation of slime protein components by electrophoresis fractionates them into high, middle and low molecular weight (MW) proteins, with high MW proteins being the most dominant^8^. Using Sanger sequencing of expressed sequence tag library of total RNA extracted from the slime gland, Haritos *et al*.^8^ obtained partial sequences of the high MW protein from *Euperipatoides rowelli*, which we hereby call ER_P1. They found ER_P1 to be a proline (Pro)-rich protein with a highly disordered structure and potential glycosylation sites^7^. Subsequently, Baer *et al*.^6^ revealed through atomic force microscopy and cryo-transmission electron microscopy observations that prior to secretion, the slime precursor consists of nanoglobular building blocks made of proteins and lipids.

**Figure 1.**
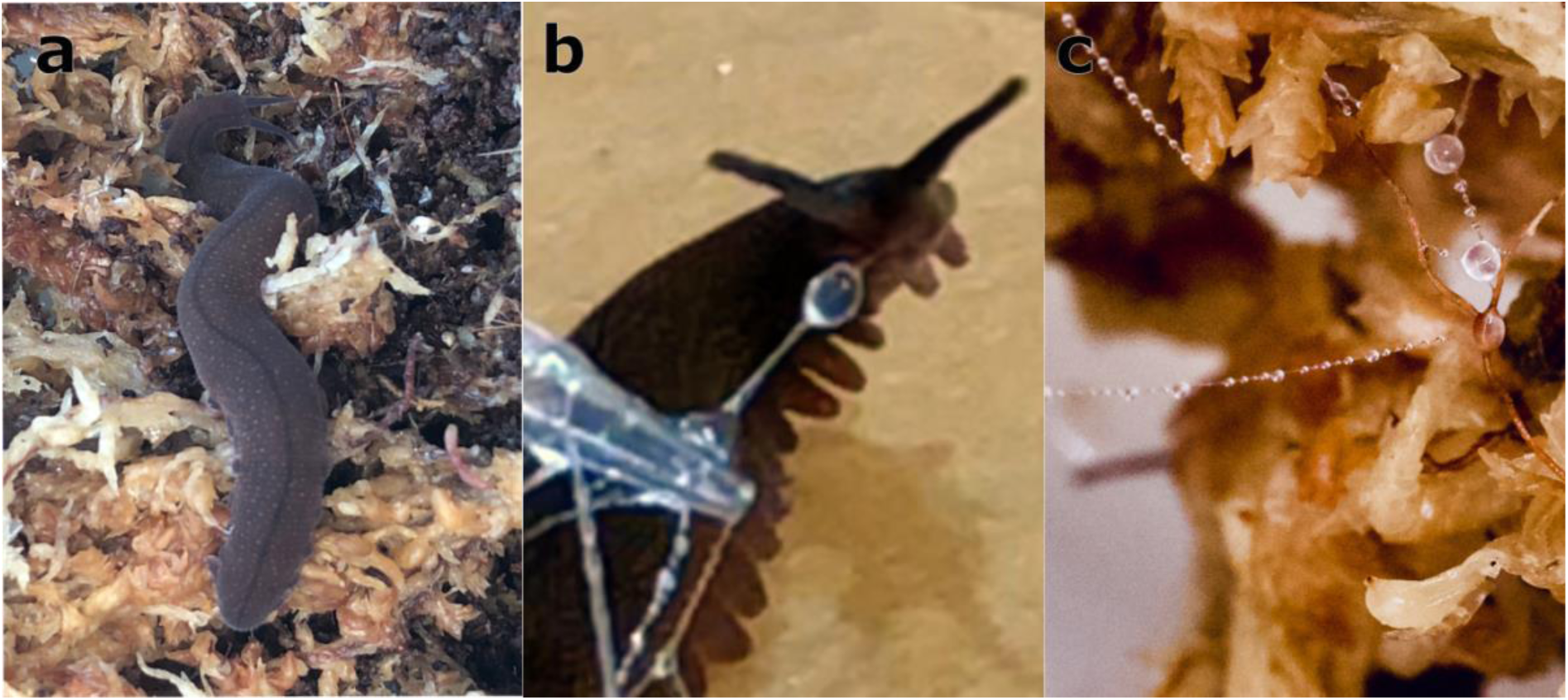
Fibrous slime of Onychophora (velvet worm). **a**. The *Eoperipatus sp*. characterized in this study. **b**. Slime ejection from the velvet worm when threatened. **c**. A close look at the slime showing droplets decorating a few threads.

Fiber formation and aqueous recycling of the slime are key physiochemical characteristics that have been investigated in recent years, but the molecular mechanisms behind fiber self-assembly are still uncertain. The most plausible explanations for intermolecular interactions governing fiber formation from the concentrated solution are lipid-protein interactions^8^, charge-charge interactions^6^, and disulfide bonding^9^. Baer *et al*.^10^ revealed that fiber formation is pH- and ionic strength-dependent. With additional prediction of phosphorylation sites on the high MW protein sequences, they proposed that the non-specific non-covalent interaction is the major driving force connecting proteins and lipids. Furthermore, crystalline β-sheet structures comprising more than 20 strands were detected in the solid slime using wide-angle X-ray diffraction^11^. β-sheet is a relatively common structural motif in adhesive materials^12^. In the velvet worm slime, it was suggested that elongated β-sheets are formed during protein unfolding triggered by shear forces, leading to rapid fiber formation and curing. The primary structure domains nucleating β-sheets are still unknown, although alternating patches with positive and negative charged residues were suggested as a possible source. The presence of divalent ions in native slime may also enhance the formation of β-sheet through phosphorylation/ Ca^2+^ interactions similar to caddisfly larvae silk ^13^. Thus, a reversible fiber assembly model combining both electrostatic interactions and shear-induced β-sheet elongation has been proposed^14^. However, experimental evidence is still lacking to fully explain fiber formation. Disulfide bonding is also a common linkage in fibrous proteins^15^ and was suggested to be related to the sticky and elastic properties of the slime^16^. However, the limited number of detected cysteine (Cys) in the partial sequence and absence of a reducing agent to reverse aggregation challenged this theory^8^.

In order to elucidate the mechanism of fiber formation, it is critical to obtain the full-length sequence of slime proteins, which has thus far eluded researchers due to the very high MW of the main slime proteins. Indeed, the aggregation of fibers is often governed by the end termini of their constitutive proteins. For example, silk fiber formation is regulated by pH-, ionic strength-, and temperature-dependent aggregation of the C- and N-terminus of silk fibroins^17,18^. To achieve this, we combined RNA-sequencing of the slime gland of an *Eoperipatus* velvet worm with high-throughput proteomic studies^19^ and successfully obtained the full-length sequences of ES_P1 and ES_P2, which are the most abundant proteins of the *Eoperipatus sp*. Further, with proteomic analyses we identified post-translational modifications (PTMs) of the slime proteins including phosphorylation of Ser, Thr and Tyr and hydroxylation of Pro. Although phosphorylation is detected, it was not identified in the high MW proteins. Instead, we find that large complexes of ES_P1, ES_P2, and smaller MW proteins are stabilized by disulfide bond. Finally, we find that the N-termini of ES_P1 and ES_P2 are highly enriched in Gly and Ser, with sequence features reminiscent of the low complexity (LC) domain of the fused in sarcoma (FUS) protein that is well-known to exhibit liquid-liquid phase separation (LLPS) ^20^. We then demonstrate with recombinant protein construct that this Gly- and Ser-rich domain also exhibits phase separation *in vitro*, suggesting that LLPS of slime protein is a central mechanism facilitating the solution-to-fiber transition during slime fabrication. Finally, we conducted solid state nuclear magnetic resonance (ssNMR) studies of ^13^C labelled slime and identified molecular entities of the slime that are located either in the flexible or the rigid domains of the slime proteins. Combined together, our results unveil new molecular insights into the velvet worm slime and its self-assembly mechanisms that provide bioinspired lessons for future synthesis of green biopolymers.

## Results and Discussion

### Phylogenetic identification

The animals, found locally in the secondary forest in Singapore, contain 21 pairs of legs with a body size ranging from 15 mm at 1 week up to 50 mm for mature animals. 12S Rrna sequences of the current specimen were compared with all known 12S rRNA sequences of Onychophora downloaded from NCBI and a phylogenetic tree on 12S rRNA was constructed with max likelihood method (**Supplementary Fig. 1**). The specimens are placed near to genus *Eoperipatus* sp. previously found in Thailand (JX568982.1).

### Full-length slime protein sequences by combined RNA-seq and proteomics

Total RNA was extracted directly from the isolated slime gland, subjected to RNAseq using Illumina Hiseq platform, and the raw sequences were assembled *de novo* using the Trinity software suite to build the transcriptome^21^. By searching the transcriptome library with Transdecoder^22^, 9,772 complete coding sequences were predicted (the assembled transcriptome and raw RNAseq data is submitted to NCBI under Bioproject PRJNA806368). Notably, we identified one transcript with very high similarity to the slime ER_P1 from the *Euperipatoides rowelli*^8^ (56% similarity and 99% alignment with E-value of 0) having a translated MW of 116 kDa, hereafter named ES_P1. The full-length sequence of the gene encoding ES_P1 was verified by PCR using cDNA as template, which led to 37 single nucleotide polymorphisms (**Supplementary Table 1**). Another transcript translating for a large MW protein (188 kDa) that shared homology with ES_P1 was also detected in the transcriptome and called ES_P2. Both proteins were characterized by a high content of Pro (17%) and Lys (10%), a limited number of Cys residues **(Fig. 2 and Supplementary Table 2)**, as well as a Gly- and Ser-rich domain in the N terminal sharing intriguing homology with the intrinsically disordered region of the FUS protein (**Supplementary Fig. 2**)^23^.

**Figure 2.**
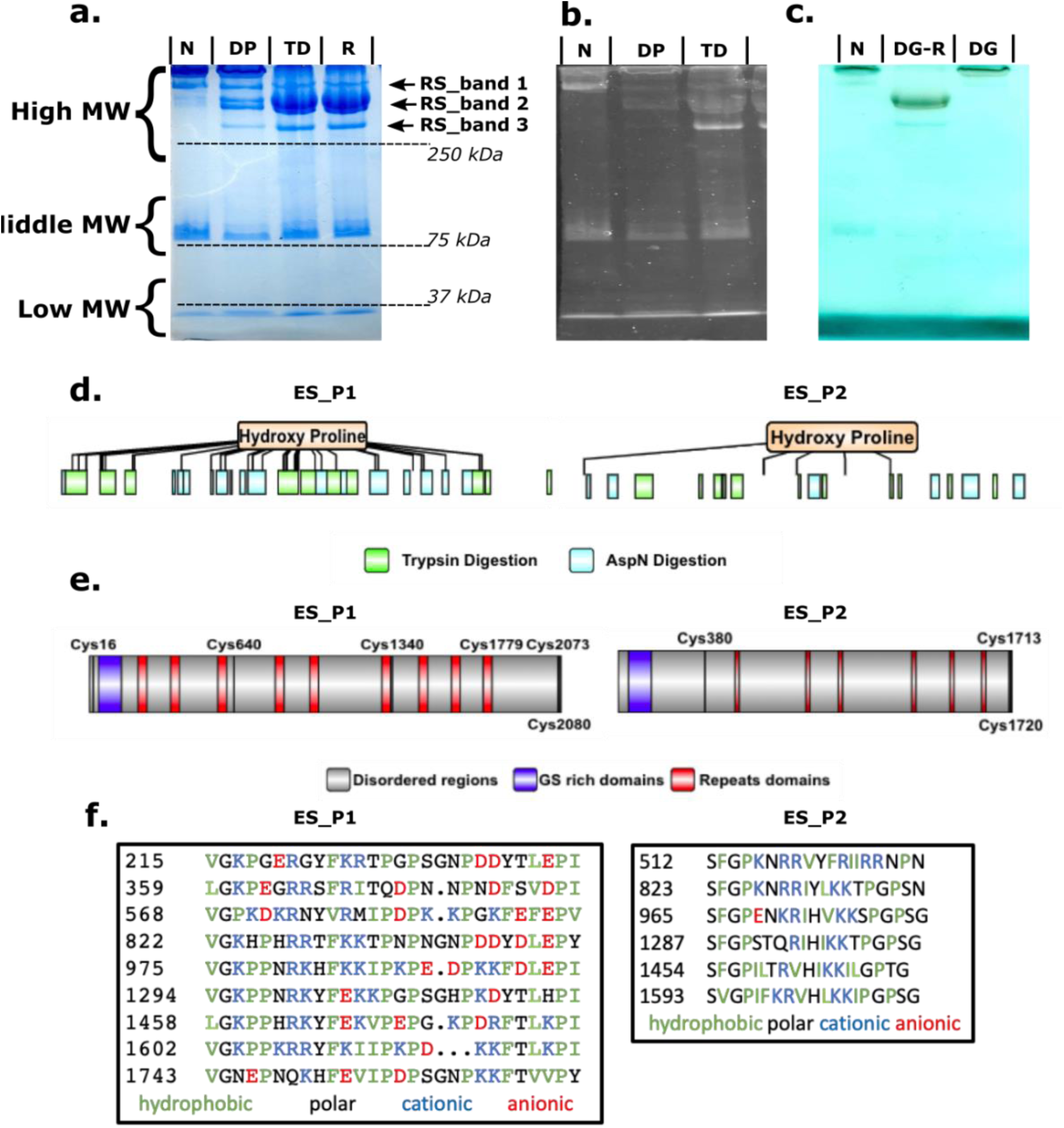
SDS-PAGE of slime proteins and full-length protein sequences of high MW slime proteins, ES_P1 and ES_P2. **a**. Long-range SDS-PAGE of the whole slime with Coomassie blue staining on native (N), dephosphorylated (DP), reduced and thermally-denatured (TD) and reduced (R) slime. **b**. ProQ stain for phosphorylated protein on N, DP, and TD slime. **c**. Alcian blue stain/silver for glycoproteins for N, deglycosylated and reduced (DG-R), and deglycosylated and non-reduced (DG) slime. **d**. Primary structure of ES_P1 and ES_P2 with peptide coverage obtained by tandem MS on fragments recovered after digestion with trypsin (green) and AspN (cyan). HyPro was also identified by LC-MS/MS for both slime proteins. **e**. Primary structures of ES_P1 and ES_P2 can be divided into 3 main domains: long predicted disordered domains (grey), Gly- and Ser-rich domains at the N-termini (purple), and repeat domains (red) located along the primary structure. The location of all Cys residues is also highlighted. **f**. Peptide sequences of repeat motifs.

To further corroborate that the identified transcripts translated into slime proteins, liquid chromatography with tandem mass spectrometry (LC-MS/MS) was conducted on native slime separated by gel electrophoresis. Slime proteins separated into high, middle and low MW regions by SDS Polyacrylamide Gel Electrophoresis (SDS-PAGE) (**Fig. 2a**). The bands from different MW regions were carefully cut off, in-gel digested, subjected to LC-MS/MS and searched against the transcriptome library of the slime gland. ES_P1 was identified as the major protein in the bands from the high MW region or reduced slime bands (RS_band 1 and RS_band_2), with 71 and 22 peptide fragments matching the translated ES_P1 transcript after trypsin and AspN digestion, respectively (**Fig. 2d**). Furthermore, 13 and 7 peptide fragments from RS_band_3 in the high MW region also matched ES_P2 after trypsin and AspN digestion, respectively.

Bands cut from the middle and low MW regions were also subjected to LC-MS/MS and probed against the slime gland transcriptome. Identified peptides matched additional transcripts with translated MWs of 71 kDa and 12 kDa, respectively. Peptides of both proteins were recovered in high abundance from the bands in their respective MW regions. The middle MW protein ES_P3 matched the middle MW proline-rich proteins from *E. rowelli* (44.6%, with 83% alignment and E-value of 4e^-141^), with a high Pro content (8.6%) in ES_P3. Similar to ES_P1 and ES_P2, a Gly- and Ser-rich domain was found at the N-terminal. The low MW protein ES_P4 did not align to any Onychophoran slime protein from the literature and its Pro content was low. In contrast, it contained a relative high amount of Cys residues (13 Cys /117 aa).

A significant feature from the LC-MS/MS data was the detection of 39 and 6 hydroxyproline (HyPro) residues (detected as oxidized Pro) in ES_P1 and ES_P2, respectively, which indicates that at least 19.6% of Pro in ES_P1 and ES_P2 are hydroxylated into HyPro. To verify the presence of HyPro in the native slime, we also conducted amino acid analysis (AAA) of hydrolyzed slime, with the HyPro retention time obtained from an external standard (**Supplementary Fig. 3**). From the overall AAA composition of the slime, we identified 4.3 +/-0.1 % of Pro and 0.38 +/-0.02 HyPro residues. Since the AAA composition measured here is for the total slime, the significantly lower content of HyPro compared with LC-MS/MS results from the individual ES_Ps suggests that HyPro residues are mostly located within the high MW proteins, a result that parallels earlier results obtained by AAA of isolated slime proteins^7^.

### Primary structure features ES_P1 and ES_P2

Prior to this study, only partial sequences of velvet worm slime proteins from *E. rowelli* had been obtained^8^. Thus, ES_P1 and ES_P2 are the first velvet slime proteins with complete and verified end-to-end primary structures. Based on homology search and structural predictions, we identify four significant features in ES_P1 and ES_P2, as highlighted in **Fig. 2f**. First, both proteins appear to be largely disordered based on predictions for intrinsically disordered proteins as discussed in more details later. Second, they contain repeat motifs about 30 amino acid long in ES_P1 and *ca*. 20 amino acid in ES_P2. Notable characteristics within the ES_P1 repeats are the presence of basic di-peptides (RR, RK, and KK motifs), a high abundance of Pro appearing as di-peptides (PP) or flanked with non-polar amino acids (IP, IIP, PI, and IP motifs), whereas anionic and polar residues appear as single residues. In ES_P2, basic di-peptides (RR, KR, KK) are also present, while Pro is usually found in the hydrophobic tripeptide PGP and FGP. In contrast to ES_P1, there are almost no anionic residues in the repeat motifs of ES_P2. Third, the N-termini of both proteins contains LC domains highly enriched in Gly and Ser (**Fig. 2f**) residues with a pattern (**G**x(0,1**)SSGGS**x**GS**-x**GGS**x(2)**SGG**x**YG**x**S**x**GGS**x(4)**GS**x**G**x(2)**G**x**S**x(5)**G**) (where x can be any amino acid) predicted with bioinformatic tools^24^. These domains share intriguing homology with LC domains of the FUS RNA-binding protein^23^, which is well-established to exhibit LLPS. Fourth, both proteins contain a small number of Cys residues (6 ES_P1 and 3 in ES_P2) that are mostly located near their extremities, namely at positions 16, 2073, and 2080 in ES_P1, and positions 1713 and 1720 in ES_P2. Since the end termini sequences of slime proteins were unknown prior to this work, these Cys residues were not detected in previous studies.

### Large MWs complex in the velvet worm slime

In gluten elastomeric proteins^25^, Cys residues located at both the C- and the N-terminus have been shown to link glutenin subunits through disulfide bonds in a tail-to-head arrangement, which stabilize and provide elasticity to the structure. A similar disulfide bridge is also known to link the heavy and light chains of *Bombyx mori* silk fibroins^26^. Based on this resemblance, we explored whether disulfide bonds could also assemble slime protein subunits into larger complexes. To this end, we enhanced electrophoretic separation of the slime proteins using long-range SDS-PAGE gels. The proteins from the native slime located in the high MW region were further separated into 4 distinct bands (band 1, 2, 3 and 4 in **Supplementary Fig. 4**). After treating the slime with DTT to reduce potential disulfide bonds, these bands shifted to the lower MW region (**Supplementary Fig. 4a**, see also **Fig. 2a**) and separated into additional distinct bands, called reduced slime band 1 (RS band 1), 2 (RS band 2, and 3 (RS band 3), confirming that proteins detected in the high MW region of SDS-PAGE gels are in fact complexes linked by disulfide bonds. To identify the components of each disulfide-linked complex in the high MW region, each band of the native slime from the long range gel was cut-off, subjected to DTT reduction, and stacked onto a second SDS-PAGE gel for further separation (**Supplementary Fig. 4b**). Proteomic analysis by LC/MS-MS of bands from this second gel showed that each band consisted of complexes of multiple proteins (**Supplementary Table 3**). Bands 1, 2, and 3 were dominated by ES_P1, whereas band 4 mostly contained ES_P2. Importantly, two additional low MW proteins, called ES_P5 and ES_P6, were detected in these complexes when the peptide fragments were probed against the translated transcriptome (**Supplementary Table 3**). These proteins were distinct from ES_P3 and ES_P4 (no significant match) identified from the middle and low MW regions of the SDS-PAGE gels of the non-reduced slime shown in **Fig. 2a**. ES_P5, with a MW of 74 kDa, was detected in all bands in high abundance. There were 16 Cys, 9.5% Ser/ 5.3% Thr evenly distributed along the primary structure of ES_P5, indicating possible sites for disulfide bond and phosphorylation, respectively. ES_P6 was only detected in band 2 and was rich in charged residues Glu (13.4%) and Lys (14.4%), often in the form of DD or KK dipeptides. Only four Cys residues were found in ES_P6: three at the N-terminus region and one at C-terminal, similar to ES_P1 and ES_P2. The relative content of Ser and Thr was similar to that of ES_P5. However, no phosphorylation modifications were detected in either ES_P5 or ES_P6.

Both phosphorylation and glycosylation staining indicated that the high MW complexes were phosphorylated and glycosylated (**Fig. 2b and 2c**). Phosphorylated residues were previously suggested to mediate nanoglobules formation^6^ or to enhance β-sheet crystalline stacking^11^, and a similar function has been identified in aquatic caddisworm silk, whereby phosphoserines of heavy fibroin interact with Ca^2+^ ions to β-rich structures and fibers^27^ via electrostatic interactions. However, phosphorylation sites in the velvet worm slime were only predicted in the high MW slime proteins and not experimentally confirmed^6^. Our Pro-Q stain after disulfide bond reduction (**Fig. 2b**) clearly indicate that phosphorylation was present mostly in the lower MW proteins of the large complexes, particularly in band 2. However, phosphorylation sites were not detected in either ES_P1 nor ES_P2 but only in peptides from a smaller MW protein (ES_P7) present in bands 1 to 3 at low abundance, at position Thr 133, Tyr 135 and Ser 136. Since phosphorylation was only detected in lower MW slime proteins, its role may be to mediate linkage of the different slime proteins in the larger complex. While our data indicate that the large MW proteins ES_P1 and P2 are not phosphorylated, we cannot rule out that additional phosphorylated sites on the smaller MW slime proteins could be detected by LC MS/MS using different enzymatic treatments.

In comparison, after disulfide bond reduction glycosylation was mainly found in the complex dominated by ES_P1 (**Fig. 2c**). To identify the carbohydrate moieties linked to the slime proteins, we conducted lectin binding assays^28^. FITC-labelled lectins showed binding to ES slime using a dot blot assay, demonstrating the presence of the specific glycans, including β-D-galactosyl(1-3)-D-N-acetyl-D-galactosamine, L-fucose, terminal α-D-galactosyl, α-D-mannose, and N-acetyl-β-D-glucosaminyl sugars (**Supplementary Fig. 5**). FITC-fluorescence based binding assay measurements indicated that α-D-mannose, and N-acetyl-β-D-glucosaminyl sugars are the most abundant glycans (**Supplementary Fig. 6**).

### Structural predictions and characterization of slime proteins

We used the AlphaFold tool^29^ for protein structure prediction, which projected that eleven and thirteen β-sheet rich domains existed in ES_P1 and ES_P2 respectively (**Fig. 3a and Supplementary Fig. 7a**), with high predicted local distance difference test (LDDT) scores, indicating correctly predicted local domains based on interatomic distances^29^. We also subjected both ES_P1 and ES_P2 to a suite of bioinformatic tools used to identify intrinsically disordered regions (IDRs), which all predicted the proteins to be largely disordered (**Fig. 3b and Supplementary Fig. 7b**). Although a high degree of disorder was confirmed, it is interesting to note that AlphaFold prediction indicates the ability of both proteins to acquire localized short secondary structures (**Figure 3c and Supplementary Fig. 7c**), corroborating crystalline β-sheets identified in the native slime by wide-angle X-ray diffraction Raman spectroscopy^11^.

**Figure 3.**
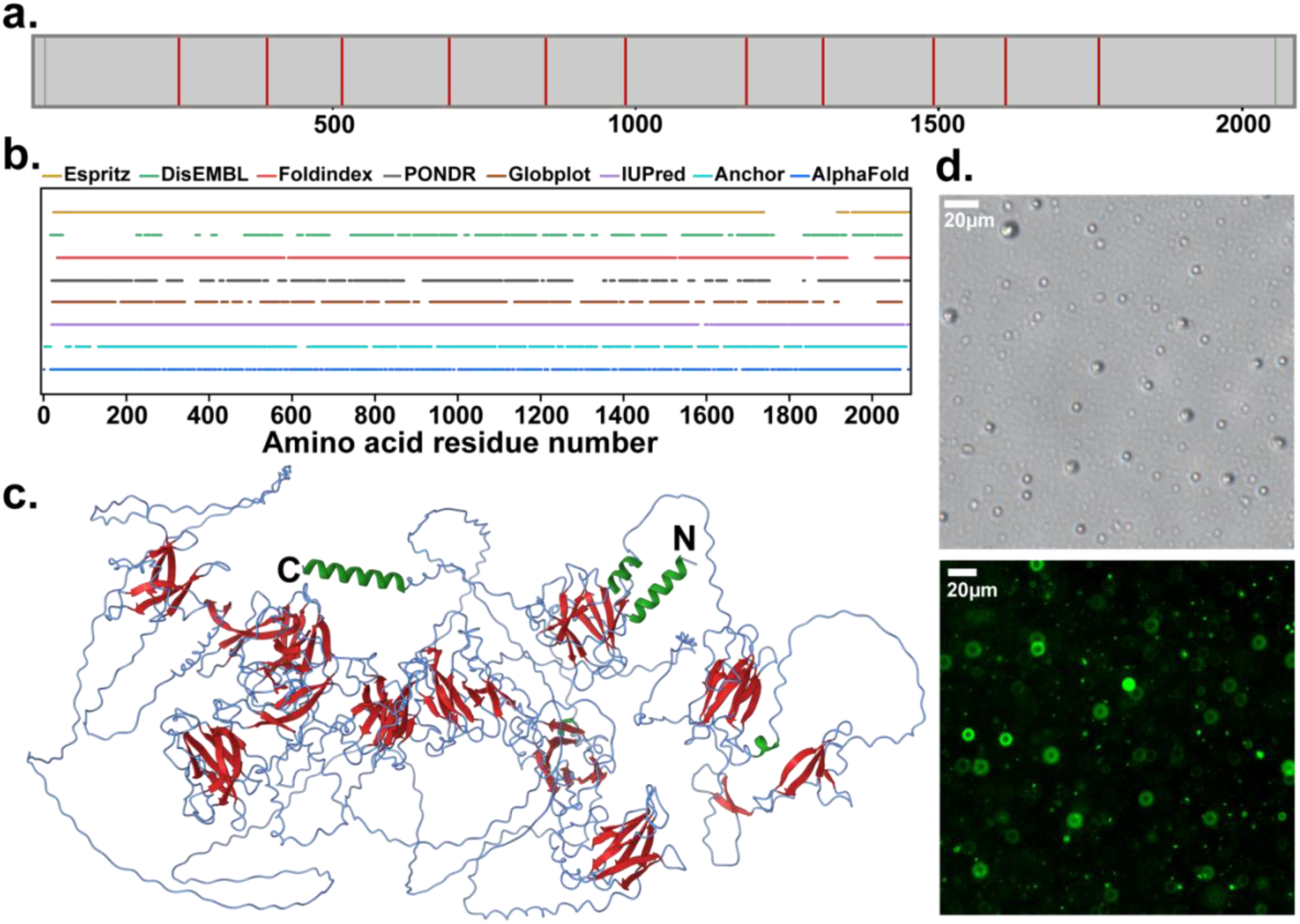
Structural predictions and LLPS of ES_P1. **a**. Secondary structure domains predicted by AlphaFold indicated as straight vertical lines along the sequence (green: α-helices; red: β-sheets). **b**. Prediction of intrinsically disordered regions within ES_P1 using various bioinformatics tools. **c**. Predicted structure of ES_P1 based on AlphaFold, with regions of secondary conformation mapped within the structure (green: α-helices, red: β-sheets). **d**. Microdroplets of recombinantly expressed N-terminus region of ES_P1 (ES_P1_31-83_, located within the purple blocks in Fig. 2e) observed by light microscopy (top), and GFP-encapsulated microdroplets of ES_P1_31-83_ observed by fluorescent microscopy (bottom).

Yet, the N-terminal regions of both proteins containing the LC domains enriched in Gly and Ser were predicted to lack any ordered structure, as evidenced by the low LDDT scores. Based on the striking resemblance of the N-termini domains with the LC domains of the FUS protein that exhibits LLPS, we hypothesized that these domains may play a similar role in ES_P1 and ES_P2, namely they may trigger the formation of concentrated droplets through LLPS. To verify this hypothesis, a construct of ES_P1 from position G31 to Y83 (referred as ES_P1_31-83_) was recombinantly expressed in *E. coli* and purified. When ES_P1_31-83_ construct solubilized in citrate-phosphate buffer (at concentration above 50 μM) was pipetted within buffer solutions at pH 4.5 to 8, microdroplets 0.4 to 5 μm in diameter spontaneously formed (**Fig. 3d**), confirming its ability to exhibit LLPS. These droplets preferably formed at 37 °C but were less visible at room temperature (**Supplementary Fig. 8**). To enhance visualization, GFP was added to the buffer solution and subsequently recruited within the droplets during phase separation as verified by fluorescence microscopy. Overall, these data suggest that the LC domain at the N-terminus in ES_P1 may promote LLPS of the slime proteins as a way to concentrate them prior to slime ejection while at the same time preventing premature aggregation. This mechanism is reminiscent of spider silk formation, although in the latter case silk spidroin precursors are concentrated through fully structured domains^30^ as opposed to IDRs in the velvet worm slime.

### Nuclear magnetic resonance (NMR) of the slime

To correlate structural and molecular features of the slime, we conducted ssNMR measurements on ^13^C-enriched slime from velvet worms. Amino acid signals with greater than 5% abundance in either ES_P1 or ES_P2 proteins were assigned to the 1D ^13^C cross polarization with magic angle spinning (CP-MAS) and direct polarization with magic angle spinning (DP-MAS) ssNMR spectra of ^13^C-enriched slime (**Fig. 4a and Supplementary Table 4**). To achieve a higher peak resolution in CP-MAS spectra as well as the acquisition of insensitive nuclei enhanced by polarization transfer (INEPT) spectra, the dried slime was wetted with a few μl of water. Peaks between 100-160 ppm can be assigned to aromatic carbons of Tyr and Phe, and Cζ of Arg, all of which consist of more than 3 mol. % in both ES_P1 and ES_P2 (**Fig. 4b**). Carboxylic peaks were not observed in the INEPT spectrum due to the absence of directly bonded protons for polarization transfer. Glu C_γ_ and Asp C_β_ peaks were more prominent in the 1D INEPT but suppressed in CP-MAS, indicating their presence in flexible regions of the proteins as well as their involvement in hydrogen bonding in the slime, which could potentially stabilize the nanoglobules in solution. Due to the overall abundance of Gly in the slime (27% predicted by AAA, **Supplementary Table 2**), the ^13^C spectrum of ^13^C-Gly labelled slime was used to identify the Gly Cα peak, and this peak was deconvoluted in the DP-MAS spectrum to identify the relative abundance of Gly in difference secondary conformations in the slime^31^. The resonance at 44.4 ppm could be assigned to Gly Cα in random coil given that the signal was significantly enhanced in the INEPT spectrum, whereas the 45.3 ppm resonance was assigned to Gly Cα in β-sheets^32^ (**Fig. 4c**). Based on the areas under the deconvoluted peaks, 16% of Gly residues were estimated to be within β-sheets while 84% of Gly were found in random conformations, allowing the latter to flexibly interact with different residues forming both inter- and intra-molecular bonds in the slime. In comparison, 5 out of 187 Gly (2.7%) and 41 out of 166 Gly residues (24.7%) in ES_P1 and ES_P2, respectively, were predicted by AlphaFold to lie within secondary structures in these two proteins (**Fig. 4d**). Next, amino acid residues within the flexible regions of the slime were assigned using 2D INEPT (**Fig. 4e**). Peaks located at 57.4, 71.8, 80.3, and 108.6 ppm in the 2D INEPT spectrum were assigned to glycan carbons C6, C2, C4 and C1, respectively, strongly indicating the presence of 6-carbon monomeric sugar moieties on the slime proteins^33^ (**Supplementary Fig. 9**). These results corroborate the presence of galactose/glucose/mannose sugars bound to proteins detected using the lectin-binding assay. Additionally, since the Ser peaks were very prominent in the 1D and 2D INEPT spectra compared to the CP-MAS spectra, we posit that glycosylation likely occurred on these Ser residues. The unassigned peak at 3.4 ppm could arise from an aliphatic group of the lipid moiety^34^. In the 2D DARR spectrum collected for 500 ms, the observed 45.3 ppm-54.0 ppm cross-peaks were assigned to Gly Cα-Lys Cα inter-residue correlations. Given that 28 pairs of consecutive GK residues exist in ES_P1 and 12 GK pairs in ES_P2, these results suggest that those pairs are located in the rigid structure of the solid slime and therefore involved in its structural arrangement (**Supplementary Fig. 10**).

**Figure 4.**
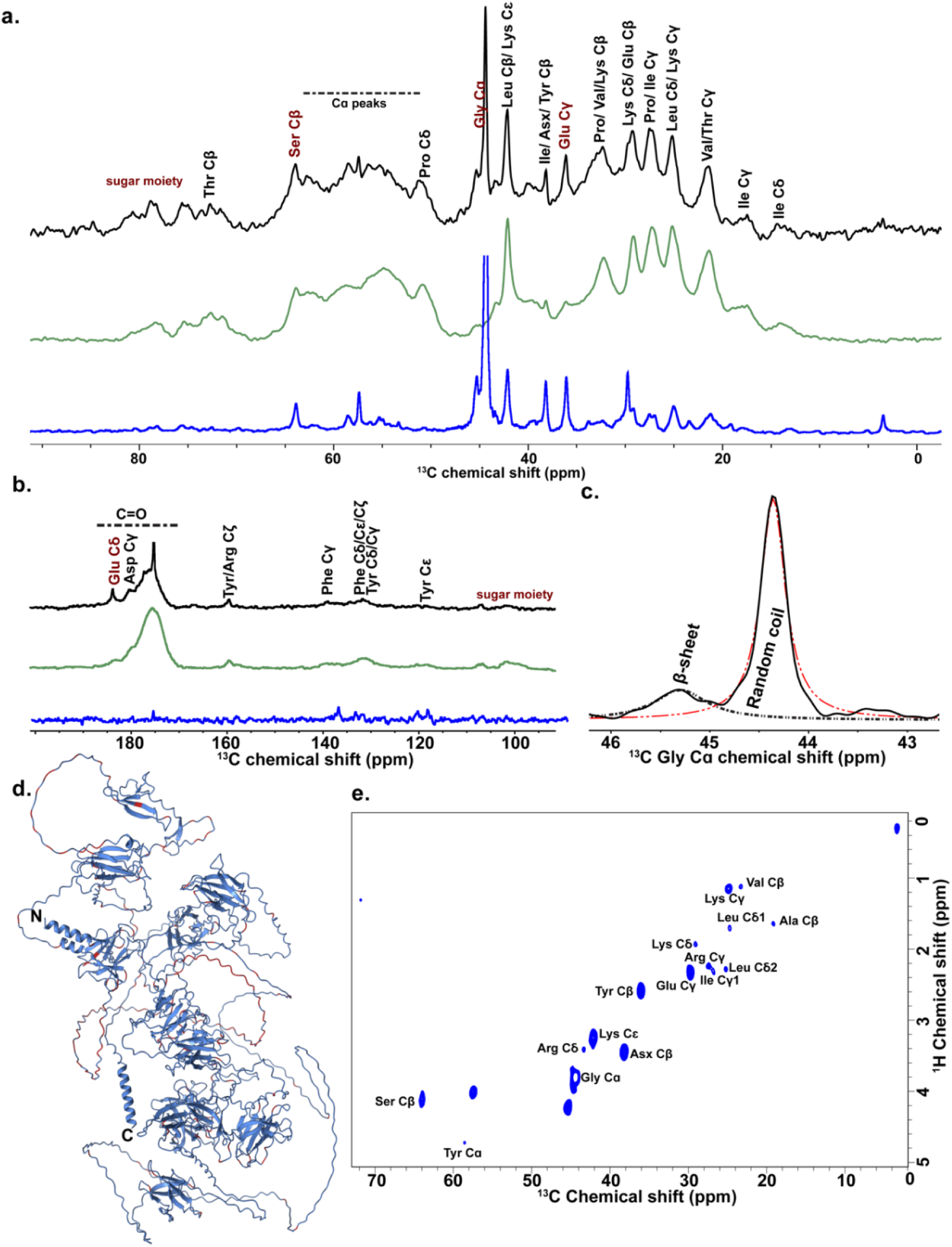
NMR characterization of the slime. **a**. 1D ^13^C ssNMR DP-MAS (black), CP-MAS (green) and INEPT (blue) spectra overlaid for *ES* slime. Residues of interest are labelled in red. **b**. Peaks beyond 100 ppm overlaid in 1D ^13^C spectra. **c**. Deconvolution of the Gly peak in DP-MAS spectrum. **d**. Mapping of Gly residues (red) on the predicted ES_P1 structure. **e**. 2D ^1^H-^13^C INEPT of hydrated *ES* slime.

### Fiber formation after thermo-mechanical treatments

From structural predictions/characterization and SDS-PAGE of reduced samples combined with LC/MS-MS, it appears that β-sheet domains and disulfide bonds are important structural and biochemical characteristics of the slime complex. To gain further insights into the role of these features on slime formation, we subjected the re-dissolved slime to thermal denaturation (to disrupt β-sheets) and/or treated it with DTT (to reduce disulfide bonds) and then attempted to draw fibers from the treated solutions. In native conditions, long fibers could readily be drawn (**Supplementary Movie 1**) as expected. When the re-dissolved slime was heat-treated to 70°C, fiber formation was inhibited (**Supplementary Movie 2**), although some weak fibers could sometimes be obtained. In contrast, incubating the slime with a high concentration of DTT (to ensure complete reduction of the disulfide bonds as verified by SDS-PAGE, see **Supplementary Fig. 11**) did not inhibit fiber formation (**Supplementary Movie 3**). Finally, the combination of both heat-treatment and DTT was the most efficient at inhibiting fiber formation, with no fibers observed in all cases (**Supplementary Movie 4**). These results indicate that thermal denaturation of the slime proteins is most efficient at preventing subsequent fiber formation.

### Updated fiber formation model

Fiber formation has recently been proposed to occur by shear-induced β-sheet unfolding followed by stabilization mediated by electrostatic interactions of phosphorylated residues^11^. An earlier suggestion was that intermolecular disulfide bonds help in stabilizing the slime upon ejection^16^. Based on our findings, we propose an updated fiber formation model in Onychophora, as illustrated in **Fig. 5**. Our results clearly support that native slime proteins do not exist as monomeric units but as multi-protein complexes linking high MW with low MW proteins by disulfide bonds through Cys residues located at the termini of either ES_P1 or ES_P2 (**Fig. 2a** and **Supplementary Fig. 11**). However, since reducing inter-molecular disulfide bonds in the resolubilized slime does not inhibit fiber drawing (**Supplementary Fig. 11** and **Supplementary Movie 3**), we conclude that that this latter mechanism is not critical for fiber formation. Multi-protein complexes linked by disulfide bridge have previously been identified in *Bombyx mori* silk fibroins consisting of heavy and light protein chains, with the linkage located at the C-terminus of the heavy protein chain^35^. Disulfide bridges may also occur in the slime proteins given that Cys residues in ES_P1 and ES_P2 are similarly placed at both termini. Interestingly, in silk fibroins, the light chain component does not directly contribute to protein structure or fiber formation but is crucial for proper cellular secretion of the heavy chain^36^. β-sheets in the heavy chains, on the other hand, are well-established to provide mechanical stability to silk fibers^37^. Disulfide-linked complexes in Onychorphora slime are discovered here for the first time, and we suggest that the low MW slime proteins may also assist in cellular secretion by preventing early aggregation, but this remains to be validated.

**Figure 5.**
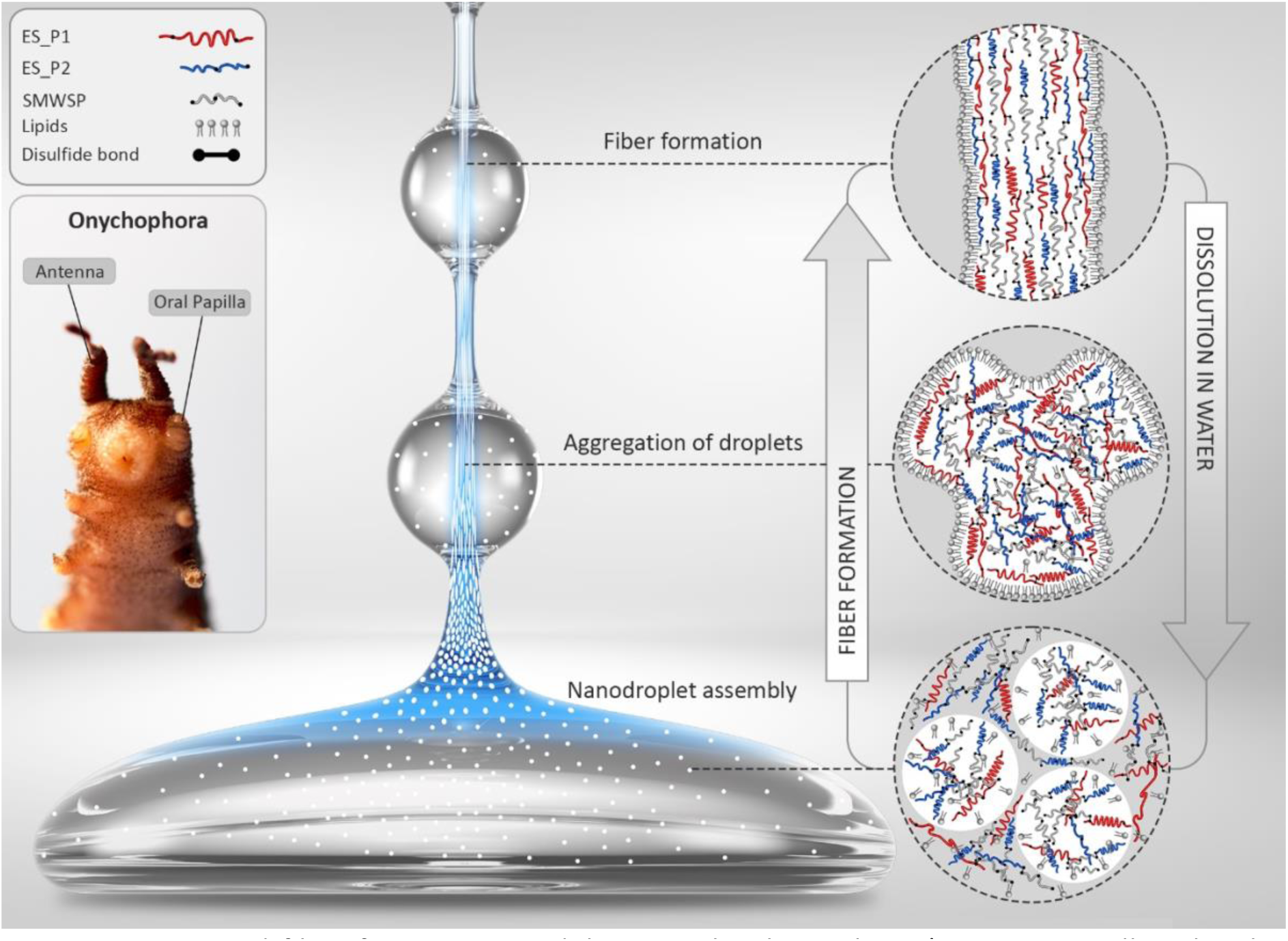
Proposed fiber formation model in Onychophora slime (SMWSP: small molecular weight slime proteins). Slime proteins are mostly concentrated within nanodroplets as disulfide-bonded complexes that also contain lipids. Protein complex and lipid phases may also be found between the droplets but at lower concentration. Upon slime ejection, β-sheets domains in ES_P1 and ES_P2 are aggregation hot spots that mediate shear-induced fiber formation and lipids migrate towards the outside of the fiber to form a hydrophobic coating (note that in the cartoon, β-sheets domains are not to scale).

Baer *et al*.^11^ suggested that β-sheets present in the initial nanodroplets undergo shear-induced unfolding, such that the content of β-sheets in the drawn fibers decreases compared to the nanodroplets. Our fiber formation experiments from the re-solubilized slime following heat treatment suggests a different mechanism. Indeed, while bioinformatic predictions clearly indicate that ES_P1 and ES_P2 are mostly in random coil configuration in the soluble state, they also confirm the presence of a few β-sheet domains that were experimentally identified in the re-dissolved slime by WAXS and FTIR^11^. The latter are thus the only structural elements that can be thermally denatured and since heat-treatment of the re-solubilized slime inhibited fiber formation, these domains appear to be critical for fiber formation. In this picture, the short predicted β-sheets domains of ES_P1 and ES_P2 (**Fig. 3b**) may act as nucleation sites for shear-induced intermolecular aggregation and fiber stabilization, concomitantly occurring with the fusion of nanoglobules into the nascent fiber.

PTMs previously detected in the slime include phosphorylation^8^ and proline hydroxylation^7^, but it was previously unknown which slime proteins are modified. According to LC MS/MS results, the large MW ES_P1 and ES_P2 are confirmed to contain HyPro whereas phosphorylation is restricted to smaller MW proteins. HyPro is the major PTM reported in spidroin proteins^38^, and has been linked to increase protein stability^39^ and mechanical properties of fibrous proteins by enhancing protein-protein interactions. It is thus tempting to suggest a similar stabilization role in slime proteins. With regard to phosphorylation, its location within a low MW slime protein is intriguing and may suggest a role in linking different proteins in the complexes, similar to the more robust disulfide bonds. We note that according to the 2D DARR NMR spectrum (**Supplementary Fig. 10**), the rigid region of the slime contains a high number (28) of the positively-charged dipeptide **GK**, most likely from ES_P1, possibly indicating electrostatic pairing with the negatively-charged phosphorylation side-chains.

In addition to proteins, the slime also contains lipids and carbohydrates. While the presence of proteins and lipids within the nanoglobules has been observed with stimulated emission depletion microscopy^6^, further evidence seems necessary to confirm their spatial distribution, especially for the proteins located outside the nanoglobules. At this stage, we assume that proteins and lipids both present inside and outside the nanoglobules. During slime ejection, we suggest that lipids move outwards and quickly coat the fibers **(Fig. 5)** possibly to enhance their hydrophobicity and prevent early disaggregation. Glycosylation is another modification whose role in slime proteins remains elusive^7^. According to our INEPT NMR spectra (**Supplementary Fig. 9**), glycosylation is located on the flexible regions of the slime, most likely on Ser residues (**Fig. 4e**), thus making it unlikely that carbohydrate side-chains play a structural role. Given the sticky nature of the slime, an adhesive functionality similar to that of the sericin in silk fibroins^40^ is more probable. However, unlike sericins that are distinct proteins from silk fibroins, glycoprotein staining on SDS-PAGE gels (**Fig. 2c**) suggests that glycosylation is post-translationally incorporated to the structural proteins ES_P1 and ES_P2.

Finally, an important finding of our study is the identification of LC sequence domains in the N-termini of both ES_P1 and ES_P2, with the construct ES_P1_31-83_ demonstrated to exhibit LLPS (**Fig. 3c**). Hence, our data suggest that the velvet worm slime may be added to the growing list of extracellular biological materials, including mussel fibers^41^, spider silk^42^, or squid beak^43^, whose biofabrication is mediated by intermediate phases formed by LLPS. Exploiting this mechanism, the velvet worm may be able to stockpile protein complexes in a concentrated state within the slime gland while at the same preventing their aggregation prior to fast ejection.

While there remain outstanding questions pertaining to the reversible liquid-to-solid transition of the viscous slime into strong adhesive fibers, the complete molecular characterization of all main slime proteins is an important step in this direction and paves the way towards sustainable fabrication of fully recyclable (bio)polymeric materials.

## Methods and materials

### Specimens and slime collection

Specimens of Onychophora were collected in the local secondary forest in Singapore (Permit No NP/RP19-037 obtained from the National Parks Board, Singapore) near the Island’s coast and maintained in the plastic boxes with perforated lids. The specimens were kept at cool temperature (20-26 °C) and sphagnum moss were used to fill 2-3 cm layer to maintain sufficient moisture. The worms were fed every week with crickets. To enrich the slime with ^13^C for NMR studies, each cricket was injected with ca. 100 μl of 5 M isotope labelled D-Glucose (U-^13^C_6_, 99% from Cambridge Isotope Laboratories) prior to feeding. The slime collection was performed every week, by stimulating the specimens with a brush and directing the ejection of slime to a pipette tip. To maximum the collection, slime was allowed to dry on the tips and stored at -20 degree until further usage. To obtain a liquid slime, the dried off slime was scratched off the tip as pieces under a dissection microscope. Slime samples are prepared as dried pieces for NMR or redissolved in minimum amount of water for other experiments.

### RNA extraction, sequencing and analysis

Total RNAs were extracted from isolated slime glands by Trizol (Thermo Fisher, United State) followed by the manufacture’s protocol. In brief, 1mL Trizol solution was added to the isolated slime gland (<50 mg). The sample was mixed by vortex for 5 minutes and further sonicated on ice with microtips, 40% power for 2-3 seconds. After 5 minutes of incubation at room temperature, 0.2 mL chloroform was added into the mixture and the mixture was subjected to centrifuge at 12,000 *g*, 4 °C for 15 minutes. The top aqueous phase was then transferred to a new tube with equal volume of 70% ethanol. Once gently mixed with the pipette, the solution was transferred to RNeasy mini kit column (Qiagen, Germany) and centrifuged at 8,000 *g* for 30 seconds to capture the RNA on the membrane. After two times wash with 700 μL RW1 solution and 600 μL RPE solution provided by the kit respectively, the spin column was transferred to a new collection column and total RNA was eluted by 30 μL RNase-free water with 2 minutes centrifuge at 8,000 *g*. The extracted RNA was stored at - 80 °C until further usage in RNA sequencing.

RNA seq libraries were then prepared using Illumina compatible NEXTflex™ Rapid Directional RNA-Seq kit according to manufacturer’s protocol. Sequencing were then performed with using 2×150 bp read length on the HiSeq 2000. The raw fastq reads were checked with FastQC^44^ and trimmed by Trimmomatic^45^. The paired end reads were *de novo* assembled by Trinity^21^ with default settings. Protein sequences were then predicted with TransDecoder^22^.

### Verification of full-length sequencing

Re-sequencing primer sets (as shown in SM) were designed in NCBI with the whole RNA seq result as database with default setting. Primers with Tm close to 65 degree with PlantiumTaq were selected for PCR. PCRs are performed according to the manufacture with adaptations. Each 50 μL reaction consisted of 250 μM dNTPs, 1x phusion plus buffer (with 1.7 mM MgCl_2_, Thermo Fisher, United State), 400 nM/L of each primer, 0.05 U Phusion plus polymerase (Thermo Fisher, United State) and 20 ng cDNA. PCR products were send for Sanger sequencing (1^st^ base, Singapore) and results were analyzed by Unipro UGENE^46^.

### Protein sample preparation & SDS-PAGE staining

Protein concentration in collected slime sample were detected in Qubit (Thermo Fisher, United State) with protein broad range assay kits. ∼100 μg were loaded into 4-15% mini gel (Biorad, England). Slime was treated as follows: (i) native slime was directly collected from the worm without further treatment; (ii) dephosphorylated (DP) slime was dephosphorylated with lambda protein phosphoatase (New England biolabs, United State) according to manufacturer’s protocol; reduced (R) and reduced with heat (RH) was carried out with 5 mM (1 mg/mL) DTT without/with heat to 70 degree for 10 minutes respectively; deglycosylated (DG) slime was treated with protein deglycosylation mix II (New England biolabs, United State) according to manufacturer’s protocol with prior denaturing. Electrophoresis was carried out at 150 V constant voltage for 60 minutes until the tracking dye reached the bottom of acrylamide gel. The gel was then incubated in sensitization buffer (30% v/v ethanol, 10% acetic acid and 10% methanol) for 1 hour. After washed with RO water, the gel was subjected to different staining chemical accordingly. Coomassie blue staining were performed according to Amini et al. ^47^ to visualize all protein contents. Pro-Q diamond phosphoprotein gel stain was performed according to the manufacture. Glyocoproteins were identified by the Alcian blue/silver staining.

PAGE on slime samples was conducted using Bio-Rad Protean II electrophoresis system by running on hand-cast 6% Bis-tris large format gels prepared at pH 6.4 with MOPS running buffer at pH 7.7 (50 mM MOPS, 50 mM Tris, 0.03% EDTA, 0.1% SDS). 15 μl of 1 mg/ml slime added to 4X NuPAGE™ LDS Sample Buffer was added per well. Electrophoresis was carried out at 150 V constant voltage initially and the current was maintained under 100 A throughout to prevent excessive heating of the PAGE system until the dye-front reached the bottom of gel. The gels were incubated in sensitization buffer containing 30% v/v ethanol, 10% v/v acetic acid and 10% v/v methanol to fix the gel and remove the excess running buffer dye, followed by quick rinse with water. Based on the sample treatment, the gels were stained using ProQ diamond stain based on manufacturer’s protocol (Invitrogen) to detect phosphoproteins, Alcian blue stain for glycoprotein detection, or Blue-silver Coomassie stain, and imaged accordingly. Ovalbumin was used as positive control for phosphorylated protein stain, and fetuin was used as control for glycosylated protein stain.

### Liquid Chromatography Tandem Mass Spectrometry (LC MS/MS)

The LC MS/MS experiments was performed according to Amini et al.^47^ with modifications. Briefly, protein bands were excised from the Coomasie-blue stained gel, and subjected to in-gel digestion with 1.25 μg trypsin (PROMEGA, sequential grade) or 1 μg Asp-N (NEB, England) for overnight digestion respectively. Digested peptides were extracted, dried and desalted with Oasis HLB 1cc 30 mg columns (Waters, WAT094225) for LC MS/MS analysis using the Easy-nLC system coupled with a Orbitrap Fusion Lumos Tribrid mass spectrometer (Thermo Scientific) and separated on a 50 cm x 75 μm Easy-Spray column. Peptides were separated over a 70 min gradient, using mobile phase A (0.1% formic acid in water) and mobile phase B (0.1% formic acid in 95% acetonitrile), and eluted at a constant flow rate of 300 nl/min using 2%-27% acetonitrile over 45 min, ramped to 50% over 15 min, then to 90% over 5 min and held for 5 min. Acquisition parameters: data dependent acquisition (DDA) with survey scan of 60,000 resolution, AGC target of 4e5, and maximum injection time (IT) of 100 ms; MS/MS collision induced dissociation in ion trap, AGC target of 1.5e4, and maximum IT of 50 ms; collision energy 35%, isolation window 1.2 m/z.

The further separated bands on long range gel were excised from zinc imidazole stain and reduced with 10mM DTT at 70 degree for 10 minutes. Individual bands were then stacked on a 2^nd^ long range gel and electrophoresis was carried out at 150 V constant voltage for another 40 minutes. After Coomassie blue staining, the protein bands from 2^nd^ long range gel were excised and in-gel digestion was performed with 1 μg Asp-N in 100 mM TEAB. Digested peptides were extracted and desalted for LC MS/MS analysis as above, but with a Orbitrap Fusion Tribrid mass spectrometer (Thermo Scientific). Separation parameters were as above, and eluted at a constant flow rate of 300 nl/min using 2%-27% acetonitrile over 45 min, ramped to 55% over 15 min, then to 95% over 5 min and held for 5 min. Acquisition parameters were as above, with isolation window of 1.6 m/z instead.

Peak lists were generated in Proteome Discoverer 2.4 (Thermo Scientific) using Mascot 2.6.1 (Matrix Science) and concatenated forward/decoy protein sequences obtained from RNAseq. Search parameters: MS precursor mass tolerance 10 ppm, MS/MS fragment mass tolerance 0.8 Da, 3 missed cleavages; static modifications: Carbamidomethyl (C); variable modifications: Acetyl (Protein N-term), Deamidated (NQ), Nitro (Y), Oxidation (M), Oxidation (P), Phospho (ST), Phospho (Y). False discovery rate estimation with 2 levels: Strict = FDR 1%, Medium = FDR 5%. Precursor mass peak (MS1) intensities were quantified by label-free quantification (LFQ) using the Minora feature detector. The peptides presenting in only one of the replicate were removed prior to quantification analysis. Total abundance of unique peptides from the same master protein was then filtered by threshold of 10^7^, and sorted to identify the major proteins in each sample/band.

### Data availability

The assembled transcriptome and raw RNAseq data is submitted to NCBI under Bioproject PRJNA806368).The raw spectra and search data were uploaded to the Jpost repository with the following accession numbers: JPST001465 (jPOST) and PXD031722 (ProteomeXchange).

### Prediction tools for disordered regions and structure

Full sequences of ES_P1 and ES_P2 were submitted to web servers to detect disordered regions from primary sequences-Espritz^48^, DisEMBL^49^, FoldIndex^50^, PONDR^51^, Globplot^52^, IUPred^53^, Anchor^54^. Secondary structures for ES_P1 and ES_P2 were predicted using AlphaFold’s google colab notebook by predictions of each protein as two halves (https://colab.research.google.com/github/sokrypton/ColabFold/blob/main/AlphaFold2.ipynb). Regions with pLDDT score below 50 were specified to be disordered. The PDB structure files were combined using ab initio domain assembly (AIDA) server (https://aida.godziklab.org/).

### Solid state Nuclear Magnetic Resonance (ssNMR)

Dried slime collected over 3 months was loaded into Bruker 3.2 mm regular wall ZrO_2_ magic angle spinning (MAS) rotor. NMR data were collected on 800 MHz Bruker Advance III instrument equipped with a 3.2 mm H/C/N EFree MAS solid-state probe. One dimensional (1D) ^1^H–^13^C cross-polarization (CP), ^13^C direct-polarization (DP), 1D and 2D ^13^C Insensitive Nuclei Enhanced Polarization Transfer (INEPT) and 2D ^13^C–^13^C Dipolar-assisted Rotational Resonance (DARR) experiments were performed with the MAS spinning frequency set at 15.151 kHz and the variable temperature set at -12 °C, which gave an actual sample temperature of 11°C based on the external calibration. Chemical shifts were referenced using the DSS scale with adamantane as a secondary standard for ^13^C (downfield signal at 40.48 ppm) and were calculated indirectly for ^1^H. The ^1^H-^13^C CP spectrum was collected with a contact time of 1200 μs. 83 kHz SPINAL-64 1H decoupling was implemented during data acquisition. 83 kHz SPINAL-64 1H decoupling was implemented during data acquisition. The recycle delays were 1.5 s and 30 s in the 1D CP and DP MAS experiments, respectively, and acquisition time was 19.1 ms in both the experiments. Additional parameters of the 2D ^13^C– ^13^C DARR experiment included 1.5 s recycle delay, 45453 Hz sweep width in both dimensions, 14.1 and 0.9 ms acquisition times in the direct and indirect dimensions, respectively. Data were processed using Topspin 3.6.

### NMR peak assignment

1D ^13^C peaks of Ile (Cβ, γ, δ at 36.1, 17.4, 14.1 ppm), Val (Cβ, γ at 32.2, 21.4 ppm), Thr (Cβ, γ at 72.6, 21.4 ppm), Leu (Cβ, δ at 42.1, 25.2 ppm), Lys (Cβ, γ, ε at 32.2, 25.2, 42.1 ppm), Glu (Cγ, δ at 36.1, 183.8 ppm), Pro (Cβ, γ, δ at 32.2, 27.2, 50.8 ppm), Ser (Cβ at 63.9 ppm), Asp (Cβ, γ at 38.2, 180.2 ppm) and Gly (Cα) were assigned based on the average chemical shifts^55^. Cδ, Cε, Cζ pf Phe, and Cγ, Cδ of Tyr are assigned to peak cluster at 129.8-132.4 ppm, Phe Cγ at 139.2 ppm, Tyr Cε at 118.2 ppm, and Cζ of Tyr/ Arg at 159.5 ppm.

### Lectin binding assay

Lectin-binding was tested using dot blot assay and fluorescence measurement using ELISA on FITC-labelled lectins^28^. Bovine Serum Albumin was used as negative control and Fetuin (WGA-binding glycoprotein) was used as positive control. For dot blot assay, nitrocellulose membrane strips were wetted with 1x Tris-buffered saline (50 mM Tris, 0.5 M NaCl, pH 7.4) with 0.05% Tween® 20 detergent (TBST) for 1 h with constant shaking. Membranes were taken out of the buffer, and 5 μl each of 0.1 mg/ml BSA, slime and fetuin were added onto the nitrocellulose membrane and the drops for left to dry for 30 min. The membranes were incubated in blocking buffer (5% w/v skim milk powder in TBST) for 2 h with continuous shaking. The membranes were washed 2x for 10 min each with TBST buffer to remove any remaining blocking buffer. The membranes were incubated in 1:200 dilution of 1 mg/ml FITC-labelled lectins in TBST buffer, and kept for continuous shaking in the dark for 2 h. The membranes were washed 2x with TBST buffer, and imaged using ChemiDoc system using Alexa fluor 488 dye settings.

For fluorescent measurements of binding assay, 50 μl triplicates each of varying concentrations of slime were added per well of transparent Nunclon™ Delta Surface 96-well plate (Thermo scientific), and left to incubate overnight at 4°C to ensure protein binding. Each well was washed twice with 100 μl 1X Phosphate-Buffered Saline (1x PBST: 137 MM NaCl, 2.7 mM KCl, 10 mM Na_2_HPO_4_, 1.8 mM KH_2_PO_4_, 0.1% w/v Tween-20) to remove unbound slime. 1 mg/ml each of FITC-labelled lectins were dissolved in PBST in 1:200 dilution to make working concentrations. 100 μl of FITC-labelled lectins were added to each well, and the plate was incubated for 2 hours in dark. Lectin solution was removed and each well was washed 1x with 100 μl PBST. 100 μl PBST was added to each well and fluorescence values were measured (ex: 490 nm; em: 525 nm). The values were plotted using Originlab.

### Recombinant purification of slime proteins

Plasmid for ES_P1_31-83_ protein was synthesized (Bio-basic Asia) and cloned into kanamycin-resistant pSUMO-LIC vector with cleavable N-terminal His-SUMO tag and expressed in chloramphenicol-resistant BL21(DE3) Rosetta T1R *E.coli* cells. Glycerol stocks were stored at –80 °C, and were used to inoculate fresh autoclaved Luria Bertani growth medium for overnight growth at 37 °C at 220rpm. These starter cultures were used to grow *E.coli* at large scale in Terrific Broth media at 37 °C until OD_600nm_ reached 1, and induced with induced with 0.5 mM isopropyl-beta-D-1-thiogalactopyranoside (IPTG) at 18°C for 18-20 h. Cells pellets were harvested by centrifugation at 6000 rpm and resuspended in lysis buffer containing 50 mM Tris–HCl, pH 7.8, 0.2 % Triton X-100, with 1 mM phenylmethylsulfonyl fluoride (PMSF) added immediately before lysis using sonicator followed by lyophilizer. Lysed sample is centrifuged at 40000 xg and the protein present in the supernatant was bound to Ni-NTA column (Cytiva) and eluted in 50 mM Tris–HCl buffer at pH 8 containing 500 mM Imidazole, 100 mM NaCl, and 1 mM TCEP. Imiadazole was removed using HiTrap column (Cytiva) before protease cleavage of the N-terminal SUMO tag. The ES_P1_(31-83)_ was collected by reverse-IMAC binding to Ni-NTA whereby the protein eluted in the flowthrough and the SUMO tag remained bound to the resin. The ES_P1_(31-83)_ protein stock for LLPS conditions was stored at 4 °C in CHAPS buffer, pH 11.

### Amino Acid Analysis

Triplicates of slime solutions in water (100 mg) were hydrolyzed in 500 mL of 6 M HCl solution containing 0.5% phenol in vacuo at 110 °C for 24 hrs. Solvents were removed by centrifugal vacuum and the hydrolysates were washed twice with water. Dried samples were kept at - 20 °C prior to analysis. For cysteine determination, triplicates of slime solutions in water were oxidized with 500 mL of performic acid (9:1 formic acid: hydrogen peroxide) in ice for 18 hrs. 100 mL of hydrobromic acid was added to the solution, after which solvents were removed by speed vacuum. Samples were then hydrolyzed in 500 mL of 6 M HCl solution with 0.5% phenol under vacuum at 110 °C for 24 hrs. Solvents were removed by speed vacuum, after which hydrolysates were washed twice with water and kept at -20 °C prior to analysis. For analysis, hydrolyzed samples (0.5 mg/ml) were dissolved in a pH 2.2 citrate buffer containing 0.1 mM Nitrotyrosine as internal standard. For performic acid-treated samples, 0.1 mM Norleucine was used as the internal standard. Composition analysis was performed with an amino acid analyzer S 433 (Sykam, Germany) using a ninhydrin buffer system.

## Acknowledgments

This research is funded by ExxonMobil through the Singapore Energy Research Center (SgEC). YTL and RMA thank the support of A*STAR Core funding and the Singapore National Research Foundation under its NRF-SIS “SingMass” scheme (RMS).

## Conflict of interest

G.P and A.U are employees of ExxonMobil who funded the project.

## References

1. Sahni, V., Harris, J., Blackledge, T. A. & Dhinojwala, A. Cobweb-weaving spiders produce different attachment discs for locomotion and prey capture. Nat Commun 3, 1106 (2012).

2. Waite, J. H. Mussel adhesion – essential footwork. J Exp Biol 220, 517–530 (2017).

3. Wolff, J. O., Grawe, I., Wirth, M., Karstedt, A. & Gorb, S. N. Spider’s super-glue: thread anchors are composite adhesives with synergistic hierarchical organization. Soft Matter 11, 2394–2403 (2015).

4. Sharma, B., Malik, P. & Jain, P. Biopolymer reinforced nanocomposites: A comprehensive review. Mater Today Commun 16, 353–363 (2018).

5. Hayashi, C. Y., Shipley, N. H. & Lewis, R. V. Hypotheses that correlate the sequence, structure, and mechanical properties of spider silk proteins. Int J Biol Macromol 24, 271–275 (1999).

6. Baer, A. et al. Mechanoresponsive lipid-protein nanoglobules facilitate reversible fibre formation in velvet worm slime. Nature Communications 1–7 (2017) doi:10.1038/s41467-017-01142-x,

7. Benkendorff, K., Beardmore, K., Gooley, A. A., Packer, N. H. & Tait, N. N. Characterisation of the slime gland secretion from the peripatus, Euperipatoides kanangrensis (Onychophora: Peripatopsidae). Comp Biochem Physiology Part B Biochem Mol Biology 124, 457–465 (1999).

8. Haritos, V. S. et al. Harnessing disorder: onychophorans use highly unstructured proteins, not silks, for prey capture. Proceedings of the Royal Society B: Biological Sciences 277, 3255–3263 (2010).

9. Röper. Analytical investigations on the defensive secretions from Perpatopsis moseleyi (Onychophora). Zeitschrift für Naturforschung 57–60 (1977).

10. Baer, A., Hänsch, S., Mayer, G., Harrington, M. J. & Schmidt, S. Reversible Supramolecular Assembly of Velvet Worm Adhesive Fibers via Electrostatic Interactions of Charged Phosphoproteins. Biomacromolecules 19, 4034–4043 (2018).

11. Baer, A. et al. Shear-Induced β-Crystallite Unfolding in Condensed Phase Nanodroplets Promotes Fiber Formation in a Biological Adhesive. ACS nano 13, 4992–5001 (2019).

12. Johnston, E. R., Miyagi, Y., Chuah, J.-A., Numata, K. & Serban, M. A. Interplay between Silk Fibroin’s Structure and Its Adhesive Properties. Acs Biomater Sci Eng 4, 2815–2824 (2018).

13. Addison, J. B. et al. Reversible Assembly of β-Sheet Nanocrystals within Caddisfly Silk. Biomacromolecules 15, 1269–1275 (2014).

14. Baer, A., Schmidt, S., Mayer, G. & Harrington, M. J. Fibers on the Fly: Multiscale Mechanisms of Fiber Formation in the Capture Slime of Velvet Worms. Integr Comp Biol 59, 1690–1699 (2019).

15. Matsuhira, T. & Osaki, S. Molecular weight of Nephila clavata spider silk. Polym J 47, 456–459 (2015).

16. Graham, L. D., Glattauer, V., Li, D., Tyler, M. J. & Ramshaw, J. A. M. The adhesive skin exudate of Notaden bennetti frogs (Anura: Limnodynastidae) has similarities to the prey capture glue of Euperipatoides sp. velvet worms (Onychophora: Peripatopsidae). Comp Biochem Physiology Part B Biochem Mol Biology 165, 250–259 (2013).

17. Hagn, F. et al. A conserved spider silk domain acts as a molecular switch that controls fibre assembly. Nature 465, 239–242 (2010).

18. Askarieh, G. et al. Self-assembly of spider silk proteins is controlled by a pH-sensitive relay. Nature 465, 236–238 (2010).

19. Guerette, P. A. et al. Accelerating the design of biomimetic materials by integrating RNA-seq with proteomics and materials science. Nature Biotechnology 31, 908–915 (2013).

20. Li, P. et al. Phase Transitions in the Assembly of Multi-Valent Signaling Proteins. Nature 483, 336–340 (2012).

21. Grabherr, M. G. et al. Trinity: reconstructing a full-length transcriptome without a genome from RNA-Seq data. Nat Biotechnol 29, 644–652 (2011).

22. Haas, B. J. et al. De novo transcript sequence reconstruction from RNA-seq using the Trinity platform for reference generation and analysis. Nat Protoc 8, 1494–1512 (2013).

23. Murray, D. T. et al. Structure of FUS Protein Fibrils and Its Relevance to Self-Assembly and Phase Separation of Low-Complexity Domains. Cell 171, 615-627.e16 (2017).

24. Jonassen, I., Collins, J. F. & Higgins, D. G. Finding flexible patterns in unaligned protein sequences. Protein Sci 4, 1587–1595 (1995).

25. Shewry, P. R., Halford, N. G., Belton, P. S. & Tatham, A. S. The structure and properties of gluten: an elastic protein from wheat grain. Philosophical Transactions Royal Soc Lond Ser B Biological Sci 357, 133–142 (2002).

26. Inoue, S. et al. Silk Fibroin of Bombyx mori Is Secreted, Assembling a High Molecular Mass Elementary Unit Consisting of H-chain, L-chain, and P25, with a 6:6:1 Molar Ratio*. J Biol Chem 275, 40517–40528 (2000).

27. Ashton, N. N. & Stewart, R. J. Aquatic caddisworm silk is solidified by environmental metal ions during the natural fiber‐spinning process. Faseb J 33, 572–583 (2019).

28. Uchiyama, N. et al. Development of a Lectin Microarray Based on an Evanescent‐Field Fluorescence Principle. Methods Enzymol 415, 341–351 (2006).

29. Jumper, J. et al. Highly accurate protein structure prediction with AlphaFold. Nature 596, 583–589 (2021).

30. Rising, A. & Johansson, J. Toward spinning artificial spider silk. Nat Chem Biol 11, 309–315 (2015).

31. Holland, G. P., Creager, M. S., Jenkins, J. E., Lewis, R. V. & Yarger, J. L. Determining Secondary Structure in Spider Dragline Silk by Carbon-Carbon Correlation Solid-State NMR Spectroscopy. J Am Chem Soc 130, 9871–9877 (2008).

32. Spera, S. & Bax, A. Empirical correlation between protein backbone conformation and C.alpha. and C.beta. 13C nuclear magnetic resonance chemical shifts. J Am Chem Soc 113, 5490–5492 (1991).

33. Arnold, A. A. et al. Identification of lipid and saccharide constituents of whole microalgal cells by 13C solid-state NMR. Biochimica Et Biophysica Acta Bba - Biomembr 1848, 369–377 (2015).

34. Knothe, G. & Nelsen, T. C. Evaluation of the 13C NMR signals of saturated carbons in some long-chain compounds. J Chem Soc Perkin Transactions 2 2019–2026 (1998) doi:10.1039/a801617h,

35. Zhou, C. et al. Silk fibroin: Structural implications of a remarkable amino acid sequence. Proteins Struct Funct Bioinform 44, 119–122 (2001).

36. Mori, K. et al. Production of a chimeric fibroin light-chain polypeptide in a fibroin secretion-deficient naked pupa mutant of the silkworm Bombyx mori. J Mol Biol 251, 217–28 (1995).

37. Keten, S., Xu, Z., Ihle, B. & Buehler, M. J. Nanoconfinement controls stiffness, strength and mechanical toughness of β-sheet crystals in silk. Nat Mater 9, 359–367 (2010).

38. Santos-Pinto, J. R. A. dos, Arcuri, H. A., Esteves, F. G., Palma, M. S. & Lubec, G. Spider silk proteome provides insight into the structural characterization of Nephila clavipes flagelliform spidroin. Sci Rep-uk 8, 14674 (2018).

39. Jr., W. G. K. PROLINE HYDROXYLATION AND GENE EXPRESSION. Annu Rev Biochem 74, 115–128 (2005).

40. Kunz, R. I., Brancalhão, R. M. C., Ribeiro, L. de F. C. & Natali, M. R. M. Silkworm Sericin: Properties and Biomedical Applications. Biomed Res Int 2016, 1–19 (2016).

41. Priemel, T. et al. Microfluidic-like fabrication of metal ion–cured bioadhesives by mussels. Science 374, 206–211 (2021).

42. Malay, A. D. et al. Spider silk self-assembly via modular liquid-liquid phase separation and nanofibrillation. Sci Adv 6, eabb6030 (2020).

43. Tan, Y. et al. Infiltration of chitin by protein coacervates defines the squid beak mechanical gradient. Nat Chem Biol 11, 488–495 (2015).

44. Andrews & S. FastQC: A Quality Control Tool for High Throughput Sequence Data. http://www.bioinformatics.babraham.ac.uk/projects/fastqc/ (2010).

45. Bolger, A. M., Lohse, M. & Usadel, B. Trimmomatic: a flexible trimmer for Illumina sequence data. Bioinformatics 30, 2114–2120 (2014).

46. Okonechnikov, K., Golosova, O., Fursov, M. & team, U. Unipro UGENE: a unified bioinformatics toolkit. Bioinformatics 28, 1166–1167 (2012).

47. Amini, S. et al. A diecast mineralization process forms the tough mantis shrimp dactyl club. Proceedings of the National Academy of Sciences 116, 8685–8692 (2019).

48. Walsh, I., Martin, A. J. M., Domenico, T. D. & Tosatto, S. C. E. ESpritz: accurate and fast prediction of protein disorder. Bioinformatics 28, 503–509 (2012).

49. Linding, R. et al. Protein Disorder Prediction Implications for Structural Proteomics. Structure 11, 1453–1459 (2003).

50. Prilusky, J. et al. FoldIndex©: a simple tool to predict whether a given protein sequence is intrinsically unfolded. Bioinformatics 21, 3435–3438 (2005).

51. Romero, P. et al. Sequence complexity of disordered protein. Proteins Struct Funct Bioinform 42, 38–48 (2001).

52. Linding, R., Russell, R. B., Neduva, V. & Gibson, T. J. GlobPlot: exploring protein sequences for globularity and disorder. Nucleic Acids Res 31, 3701–3708 (2003).

53. Erdős, G., Pajkos, M. & Dosztányi, Z. IUPred3: prediction of protein disorder enhanced with unambiguous experimental annotation and visualization of evolutionary conservation. Nucleic Acids Res 49, gkab408. (2021).

54. Dosztányi, Z., Mészáros, B. & Simon, I. ANCHOR: web server for predicting protein binding regions in disordered proteins. Bioinformatics 25, 2745–2746 (2009).

55. Wishart, D. S., Bigam, C. G., Holm, A., Hodges, R. S. & Sykes, B. D. 1H, 13C and 15N random coil NMR chemical shifts of the common amino acids. I. Investigations of nearest-neighbor effects. J Biomol Nmr 5, 67–81 (1995).

