## Supplementary Materials for "Complete sequences of the velvet worm slime proteins reveal that slime formation is enabled by disulfide bonds and intrinsically disordered regions"

#### Supplementary Tables

**Supplementary Table 1.** Single polymorphism as capital letter of ES_P1 and ES_P2 detected in Re-sequencing with designed primers comparing to the sequences obtained from RNAseq and the related changes in amino acid composition.

| | Position of RNAseq | RNAseq | Amino acid | Re-seq with primers | Amino acid |
|---|---|---|---|---|---|
| ES_P1 | 73 | aGa | Arg | aCa | Thr |
| | 409 | cCt | Pro | cTt | Leu |
| | 447 | Ctt | Leu | Ttt | Phe |
| | 519 | Gta | Val | Ata | Ile |
| | 527 | acA | Thr | acT | Thr |
| | 701 | ccT | Pro | ccC | Pro |
| | 790 | aCt | Thr | aTt | Ile |
| | 893 | gtC | Val | gtT | Val |
| | 969 | Cgt | Arg | Ggt | Gly |
| | 1517 | ggC | Gly | ggA | Gly |
| | 1975 | aGa | Arg | aAa | Lys |
| | 2734 | aAa | Lys | aGa | Arg |
| | 2795 | aaA | Lys | aaC | Asn |
| | 2936 | ccA | Pro | ccT | Pro |
| | 2937 | Aat | Asn | Cat | His |
| | 2951 | ttC | Phe | ttT | Phe |
| | 2952 | AAA | Lys | Gaa | Glu |
| | 2959 | aTa | Ile | aGa | Arg |
| | 2963 | ccA | Pro | ccT | Pro |
| | 2995 | gAA | Glu | gTG | Val |
| | 2999 | ccA | Pro | ccT | Pro |
| | 3002 | atA | Ile | atT | Ile |
| | 3008 | ccT | Pro | ccA | Pro |
| | 3011 | gaT | Asp | gaA | Glu |
| | 3012 | Gat | Asp | Aat | Asn |
| | 3023 | gaG | Glu | gaT | Asp |
| | 3024 | CCa | Pro | GAa | Glu |
| | 3032 | gAC | Asp | gaT | Asp |
| | 3038 | gaA | Glu | gaT | Asp |
| | 4028 | ggG | Gly | ggA | Gly |
| | 4091 | aaC | Asn | aaT | Asn |
| | 4203 | Cca | Pro | Gca | Ala |
| | 4784 | AAA | Lys | aaG | Lys |
| | 5234 | aaT | Asn | aaA | Lys |
| | 5850 | Ata | Ile | Gta | Val |

|  |  |  |  |  |  |
|---|---|---|---|---|---|
|  | 5914 | gCg | Ala | gTg | Val |
|  | 5923 | aCt | Thr | aTt | Ile |
| **ES_P2** | 152 | Gat | Asp | Aat | Asn |
|  | 1650 | cAt | His | cGt | Arg |

**Supplementary Table 2**. Amino acid analysis of slime comparing sequences ES_P1 and ES_P2.

| Amino acid | % in ES slime | ES_P1 | ES_P2 |
|---|---|---|---|
| Aspartate/Asparagine | 8.8 ± 0.2 | 11.3 | 12.3 |
| Glutamate/Glutamine | 11.5 ± 0.0 | 10.1 | 8.7 |
| Threonine | 3.9 ± 0.2 | 6.7 | 4.1 |
| Serine | 4.7 ± 0.7 | 5.2 | 6.7 |
| Tyrosine | 2.6 ± 0.4 | 3.4 | 3.9 |
| Phenylalanine | 1.9 ± 0.2 | 3.4 | 3 |
| Proline | 4.3 ± 0.1 | 17.7 | 16 |
| Hydroxyproline | 0.4 ± 0.0 | 3.5 | NA |
| Glycine | 27.1 ± 0.8 | 9 | 9.6 |
| Cysteine | 1.9 ± 0.1 | 0.3 | 0.2 |
| Alanine | 4.8 ± 0.2 | 1.9 | 2.6 |
| Valine | 4.3 ± 0.5 | 5.6 | 5.6 |
| Methionine | 1.0 ± 0.2 | 0.6 | 0.7 |
| Isoleucine | 2.7 ± 0.1 | 3.7 | 6 |
| Leucine | 4.3 ± 0.1 | 3.2 | 5.7 |
| Histidine | 3.8 ± 0.3 | 2.9 | 1.6 |
| Lysine | 8.5 ± 0.2 | 10.2 | 9.2 |
| Arginine | 3.6 ± 0.2 | 4.8 | 3.6 |

**Supplementary Table 3**. Proteins detected in different complex from LC MS/MS.

| Position in native gel | Proteins detected |
|---|---|
| Band_1 | ES_P1, ES_P5 |
| Band_2 | ES_P1, ES_P5, ES_P6 |
| Band_3 | ES_P1, ES_P5 |
| Band_4 | ES_P2, ES_P5 |

**Supplementary Table 4.** $^{13}C$ peak shifts in ppm assigned for amino acid residues in *Eoperipatus sp.* slime based on the average chemical shifts.

| Amino acid | Cβ | Cγ | Cδ | Cε | Cζ |
|---|---|---|---|---|---|
| Ile | 36.1 | 17.4 | 14.1 |  |  |
| Val | 32.2 | 21.4 |  |  |  |
| Thr | 72.6 | 21.4 |  |  |  |

<table>
<tr><td>Leu</td><td>42.1</td><td></td><td>25.2</td><td></td><td></td></tr>
<tr><td>Lys</td><td>32.2</td><td>25.2</td><td></td><td>42.1</td><td></td></tr>
<tr><td>Glu</td><td></td><td>36.1</td><td>183.8</td><td></td><td></td></tr>
<tr><td>Pro</td><td>32.2</td><td>27.2</td><td>50.8</td><td></td><td></td></tr>
<tr><td>Ser</td><td>63.9</td><td></td><td></td><td></td><td></td></tr>
<tr><td>Asp</td><td>38.2</td><td>180.2</td><td></td><td></td><td></td></tr>
<tr><td>Phe</td><td></td><td>139.2</td><td colspan="3">129.8-132.4</td></tr>
<tr><td>Tyr</td><td></td><td colspan="2">129.8-132.4</td><td>118.2</td><td rowspan="2">159.5</td></tr>
<tr><td>Arg</td><td></td><td></td><td></td><td></td></tr>
</table>

### Supplementary Figures

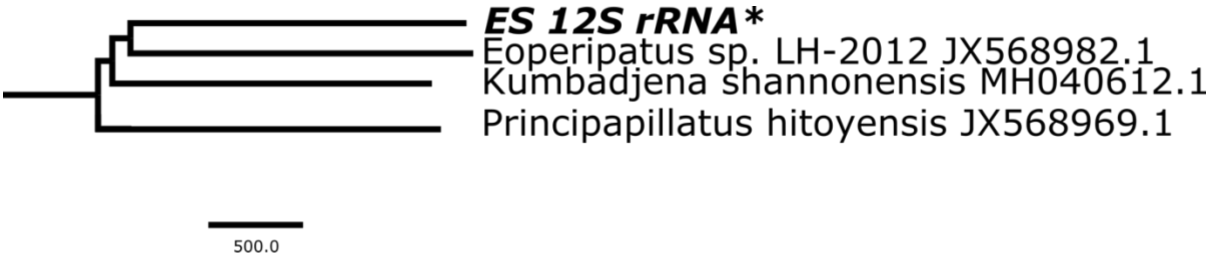


**Supplementary Figure 1.** A partial phylogenetic tree indicating the position of ES 12S rRNA in the whole phylum of Onychophora.

```
FUS-LC   MASNDYTQQATQSYGAYPTQPGQGYSQQSSQPYGQQSYSGYSQSTDTSGYGQSSYSSYGQ 60
ES_P1-NT MK-----ILLSVLVLLIVVEC--GNSRKI-RH-------------------RGGSR---R 30
ES_P2-NT ME-----MMYTLFFLLFGIVHGQGDGWVL-QP-------------------DGSYMSYGD 35
         *         :            * .    :                     ..      

FUS-LC   SQNTGYGTQSTPQGYGSTGGYGSSQSSQSSYGQQSSYPGYGQQPAPSSTSGSYGSSSQS- 119
ES_P1-NT --------GSGGGSGGSSGGS--SGG----------------------SDGSYGGSDGGS 58
ES_P2-NT --------GSSGGSYGSTGGS--YDG----------------------SGGLYGGSSGGS 63
                  *   . **:**     .                      :.* **.*. . 

FUS-LC   -SSYGQPQSGSYSQQPSYGGQQQSYGQQQSYNPPQGYGQQNQYNSSSGGGGGGGGGNYG 178
ES_P1-NT GGSYGDSGGGSGDSTGSNGG------------------PGDSYSESGGSSGDGGSGGSYG 100
ES_P2-NT ---YGELGSGG------------------------------LFGGS--GGGGFGPGGSYG 88
            **:  .*.                               :. *  ..*. * **.**

FUS-LC   QDQSSMS-----------------------SGGGSGGGYGNQDQSGGGGSGGYGQQDRG- 214
ES_P1-NT GSDGGPGGSYGG-------------------SGGSGGGGGGGGGGGGGGSGSDNNPPEGY 141
ES_P2-NT GFDGGLGGSSGGSQGLPGNGWILQPDGSYLKYEGSG--GGGGGGGGGGGSGSDGPPGNGW 146
           :.. .                          ***   *. . .******. .   .* 
```

**Supplementary Figure 2.** Sequence alignment for FUS-LC domain with N-terminal sequence of ES_P1 and ES_P2 proteins.

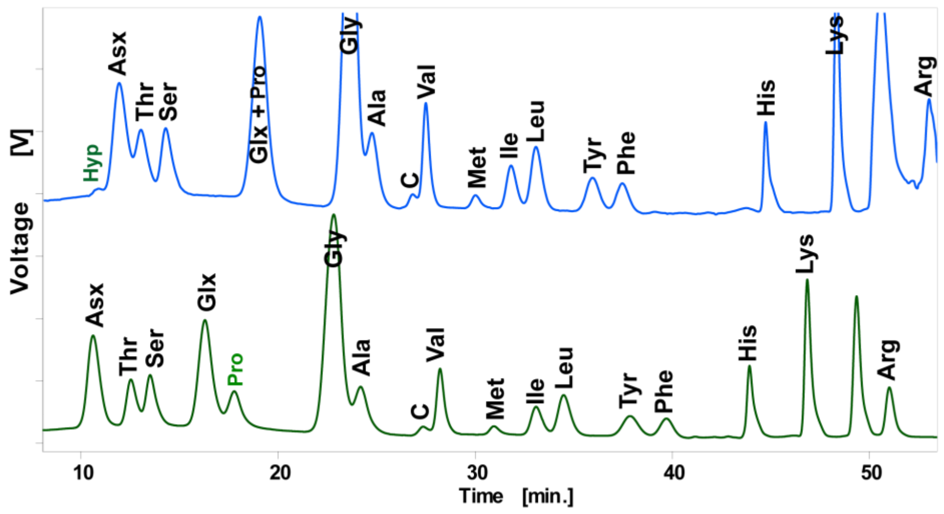


**Supplementary Figure 3**. Amino acid chromatogram of the native slime.

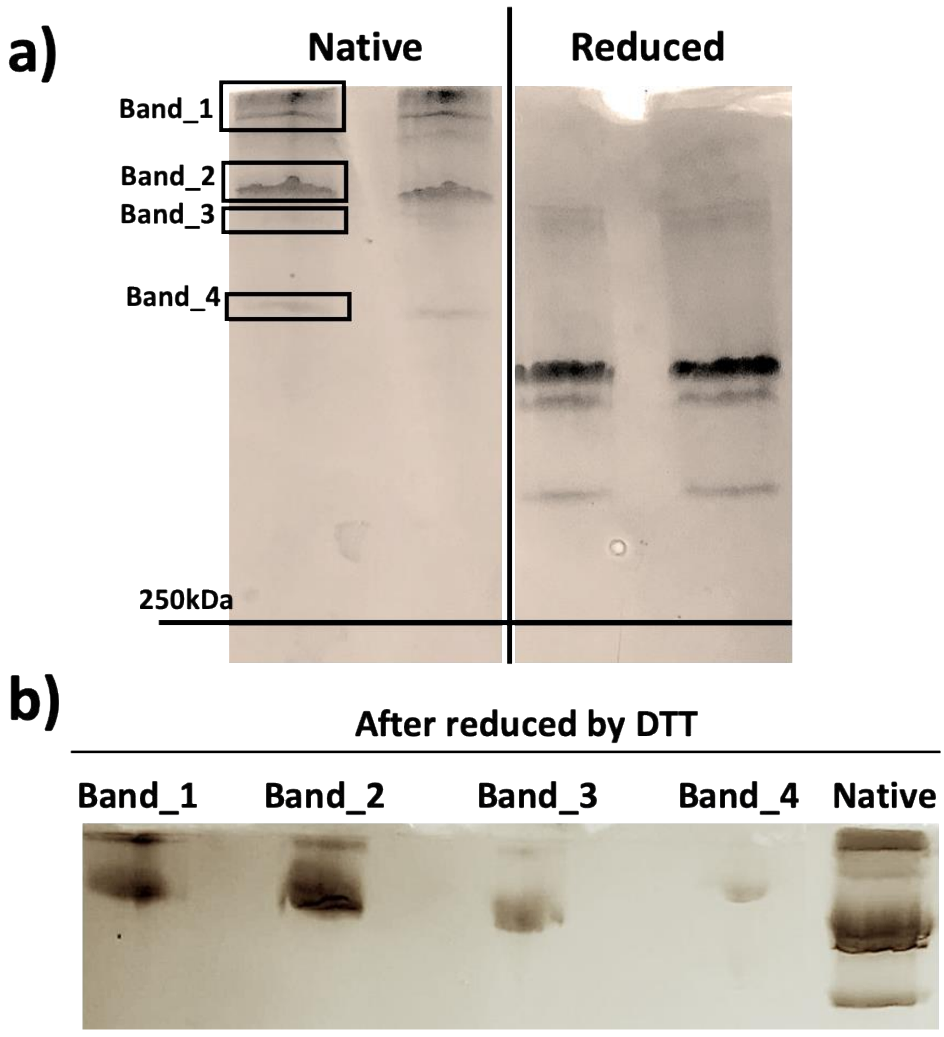


**Supplementary Figure 4.** Long range gel of native and reduced slime by DTT in duplicate a and second SDS-page on 4 bands cut off from native proteins in the long range gel, subjected to reducing disulphide bonds by DTT respectively b. All bands from second SDS-page were cut off for LC MS/MS analysis.

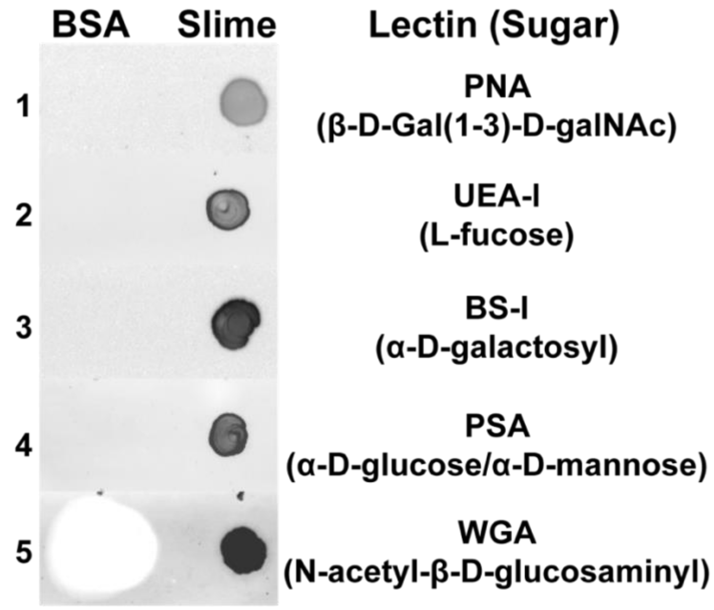


**Supplementary Figure 5.** Dot plot assay for detection of lectins binding to slime. Peanut agglutinin PNA, *Ulex europaeus* agglutinin UEA-I, *Bandeiraea simplicifolia* lectin-IBS-I, *Pisum sativum* lectin PSA, and wheat germ agglutinin WGAwere applied to ES slime.

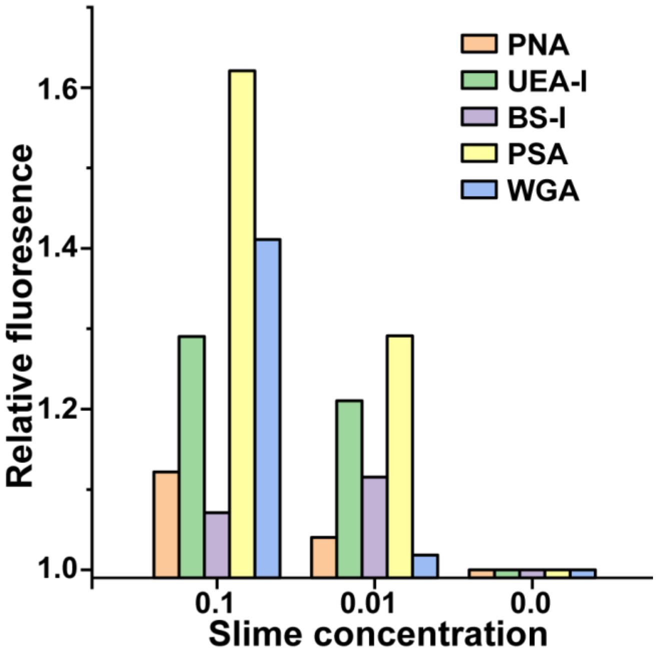


**Supplementary Figure 6.** Fluorescence measurements for detection of lectins binding to slime. Peanut agglutinin PNA, *Ulex europaeus* agglutinin UEA-I, *Bandeiraea simplicifolia* lectin-IBS-I, *Pisum sativum* lectin PSA, and wheat germ agglutinin WGA were applied in the lectin assay to ES slime. PSA and WGA bound more strongly to increasing concentrations of slime.

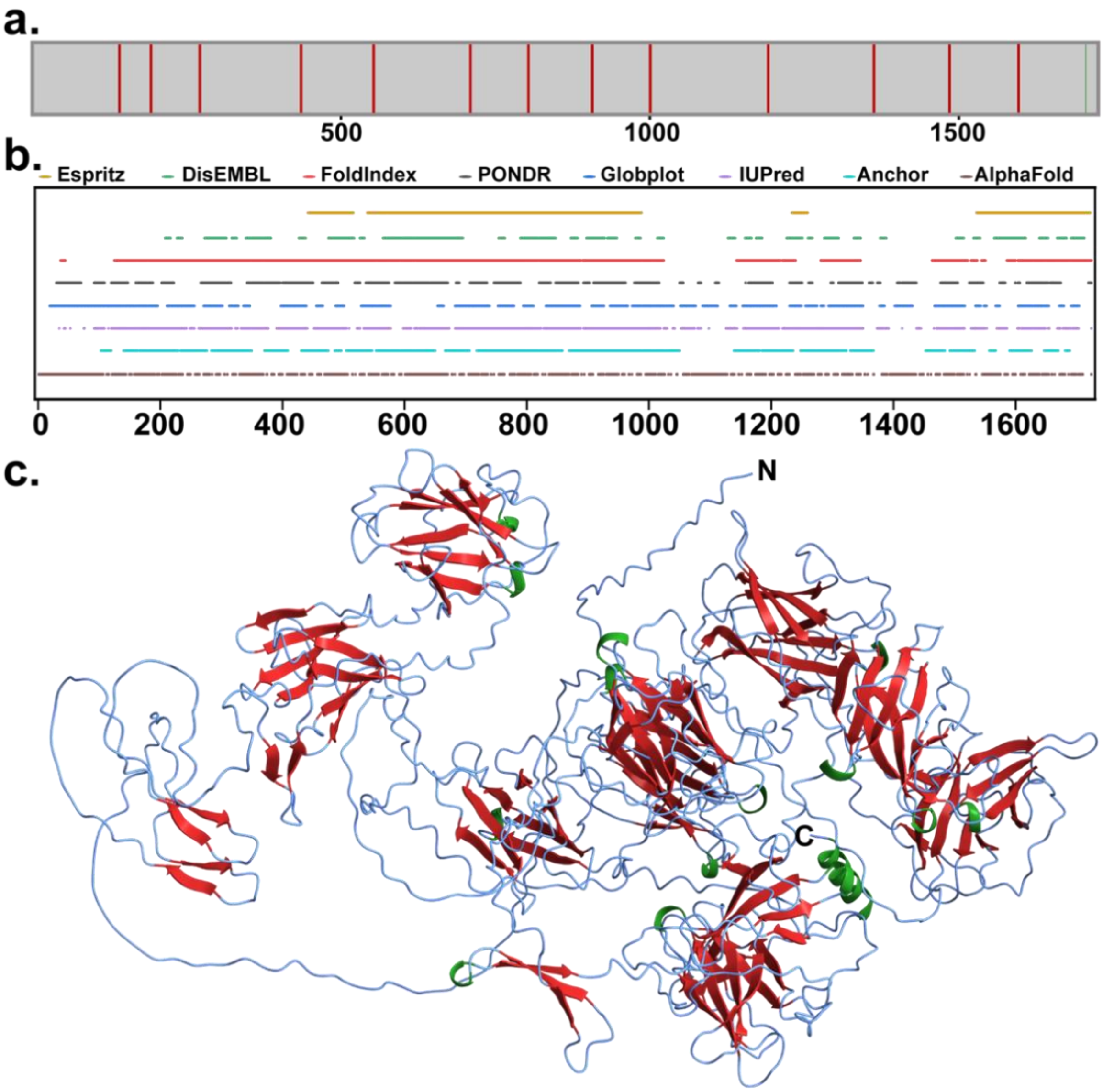


**Supplementary Figure 7.** Structural predictions of ES_P2. **a**. Prediction of secondary structure domains predicted by AlphaFold indicated as straight line along the sequence top, green: α-helix, red: β-sheet b. intrinsically disordered regions within ES_P1 using bioinformatics tools and c. Predicted structure of ES_P1 based on AlphaFold, with regions of secondary conformation mapped within the structure green: α-helix, red: β-sheet.

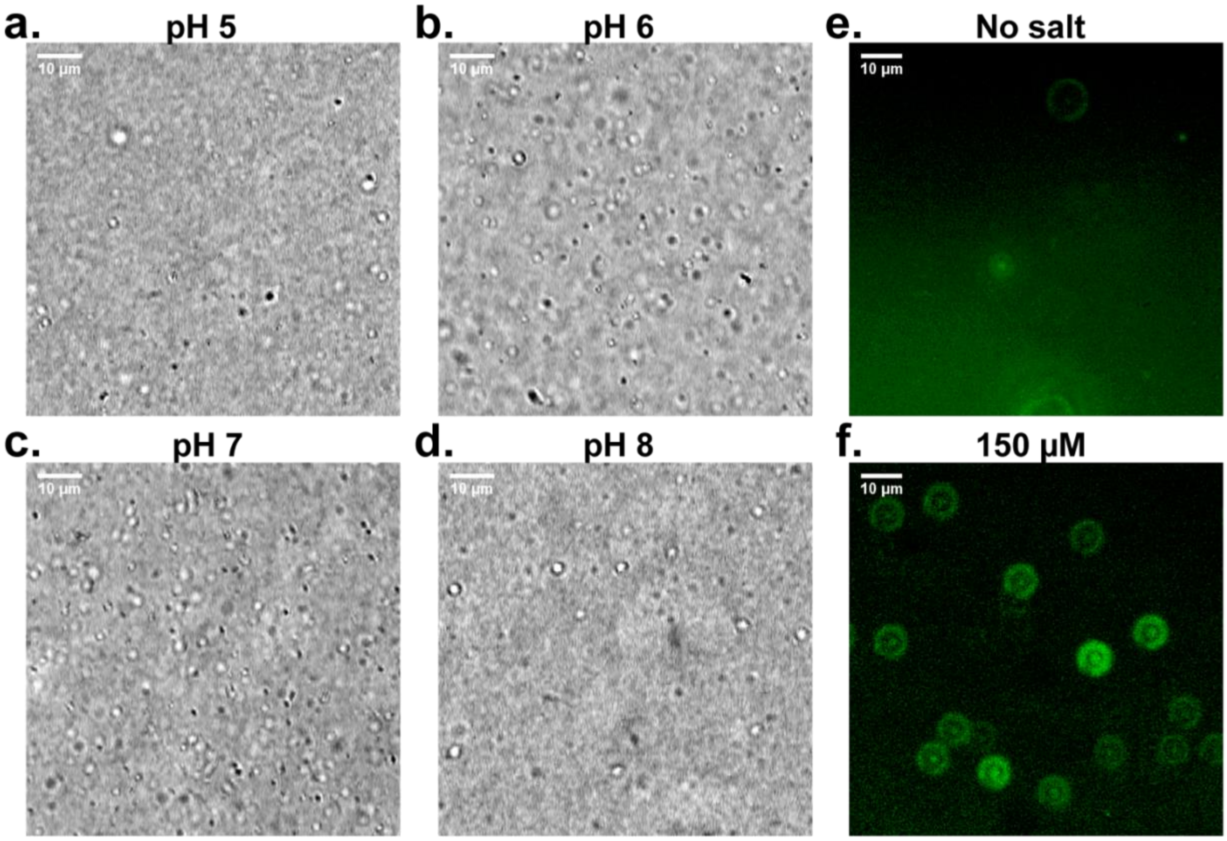


**Supplementary Figure 8. a-d.** Microdroplets of 60 µM ES_P1$_{31\text{-}83}$ recombinant protein observed under optical microscope at pH 5 a ,6 b ,7 c, and 8 d. **e-f.** GFP-encapsulation within ES_P1$_{31\text{-}83}$ droplets observed under fluorescence microscope in the absence e and presence f of salt at 150 µM concentration.

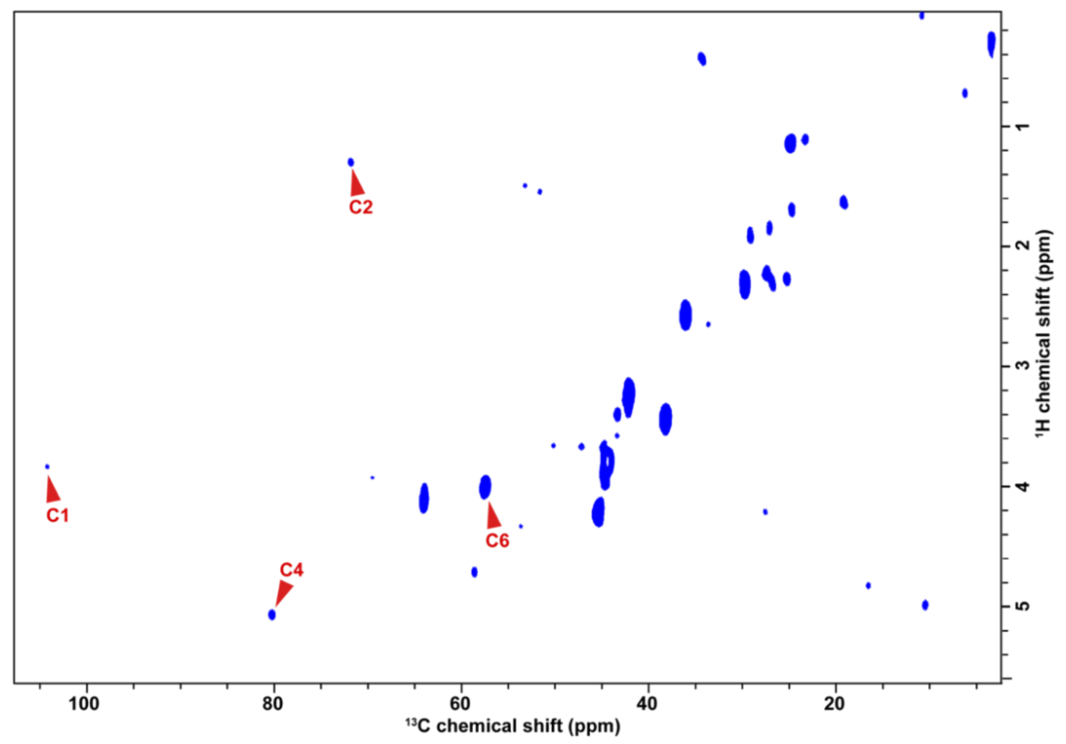


**Supplementary Figure 9**. Zoomed-out of the 2D INEPT spectrum indicating carbohydrate peaks with $^{1}H$ and $^{13}C$ chemical shifts within 60-110 ppm.

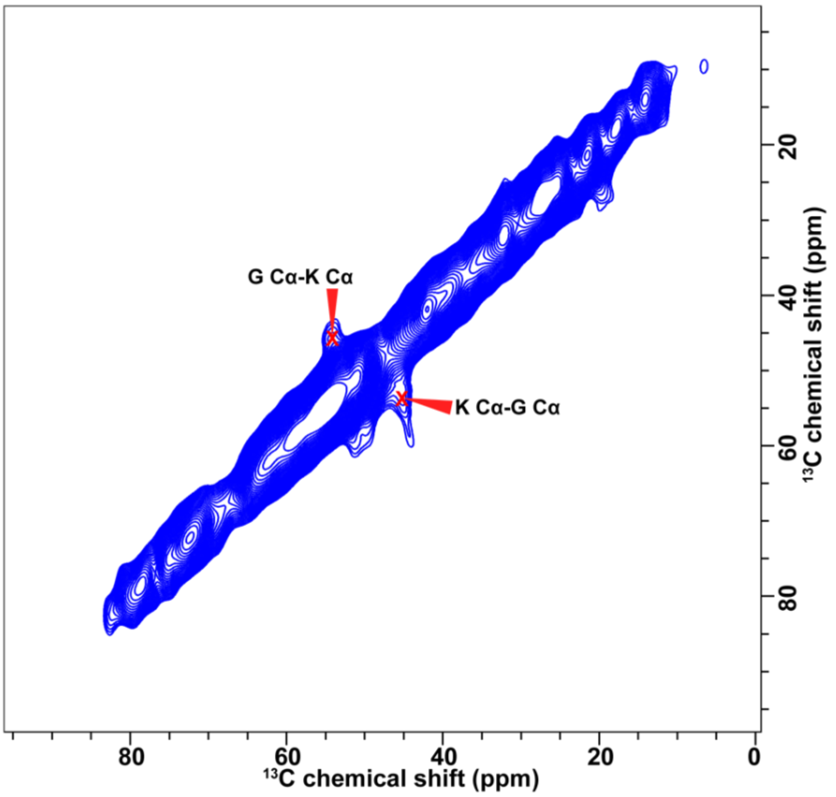


**Supplementary Figure 10.** 2D $^{13}C$-$^{13}C$ DARR spectrum with 500 ms mixing time showing G/K correlation cross-peaks along the diagonal.

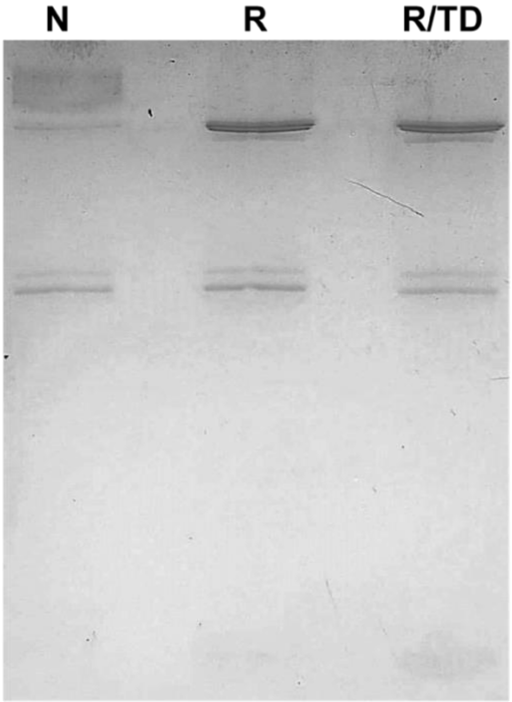


**Supplementary Figure 11.** Complete reduction of disulphide bond at high DTT concentration. The gel lanes indicate native slime N, slime reduced with DTT R, and reduced slime followed by thermal denaturation R/TD.

#### Supplementary Movies

**Movie 1**. Fiber drawn from native slime.
**Movie 2**. Heat inhibits fiber formation.
**Movie 3.** Reduced disulphide bond does not inhibit fiber formation.
**Movie 4.** Combination of heat and reduced disulphide bond inhibit fiber formation.

#### Supplementary Data

#### Protein Sequences:

**>ES_P1 (hydroxyprolines are highlighted in grey).**
MKILLSVLVLLIVVECGNSRKIRHRGGSRRGSGGGSGGSSGGSSGGSDGSYGGSDGGSGGSYGDSGGGSGDST
GSNGGPGDSYSESGGSSGDGGSGGSYGGSDGGPGGSYGGSGGSGGGGGGGGGGGGGSGSDNNPPEGY
YDPNKSGPLPPGFEAPPGYEGGEWHPGPDGTMVRTIVEEEPGTETTELVPPDYDPQITPVGQPGSPGYHDIIG
VGKPGERGYFKRTPGPSGNPDDYTLEPIKSPENPEPVDPNDPNAPRVIRTKKNKNPTLQVGNKDNPQYFGLKP
DPKNPGHFTLVPRKMHPGKPGHRKPGHPKGGKGKPGRGKPGHGKPGDGGALSPEEYEKLKQNPVMKYPVF
TLGKPEHRRSFRITQDPNNPNDFSVDPIGNENDPQSPADPNDPEAPEIIPQGEKGKPDYNVVIGVNPKKKRDY
VQMKPDPKNPQQFNFDPIFMEPQQEEPQQEQPTPAGGEPEVFLDPGSESVTPEEYERLKQNPKIDRKKGSPIL
HLGYEPYRRPFRIKQNRRNPLDFDVQPIGSDNDPNSKPDKDDPYKPVVYPLGEKGQPDYSQIIGVGPKDKRNY
VRMIPDPKKPGKFEFEPVLLKPGEPIPFVTETEPPPTPTPEPTVPPPPPTHPPTTKKPEEEEPCFEDLPEDPEFRRV
RHPGRRRPYQIISLGPKDHRQHFKKTPKSNNPDDYDIEPYDPDTPDHKPKADDPYAPEVIPQGQPGTPSFEPIL
MTGPKDNRKPYKLVHHPENPSRSSFVPVRPLPKKPGQKHPKFVNAPRQIRPRKKDEPRQPDDEPEEGEPEFTD

LPDRPRFKHVKQPKRPYTLISVGKHPHRRTFKKTPNPNGNPDDYDLEPYDEDTPDHKPNPHDPSVPKVHKNG
KKGTPDYQPVIEIGPINKRKLYKIKPHPSNPNKVEFVPVKKIMRKGKPFFKELPRRTATKRFRPRTTPKPNEDDDV
PDDLPYDPEVRPHRTPGQKYPTQIISVGKPPNRKHFKKIPKPEDPKKFDLEPIDPDDPDEPLDPEDPNTPKVYPN
PEGNPVIETGTPDDPKHHEVNPDPEDPEDPERVSFTPVKRLGEPGEKNPKFEKIPRKSHPFHTKRPEEATAPEPE
ENEPVFEDIPYNPTFKTLHRKRPDRPAQIIALGRPPHRQEFIKEPGRSGKPHDFKLSPYDSRRPDKKPDPNNPNK
PEVYPARGKPGKPGYQPPVIKTGPKDDPEYFEIHPDPSDPKNPNKVLFVPVKARPGKPGTRPKFSPKEPRRYHP
YHDPLKPRPTTEEPEDGTEYFVDQPEDPEVTTLFRPLRKYPSQLIAVGKPPNRKYFEKKPGPSGHPKDYTLHPID
PKHPDKSLNPKDSKTPKCFPGEGKPGTPNYKPPVIQTGSPDNPKFYRVIPNPKDPNNPESVEFIPVKKDKKNPK
KFKPLPRNVRIYHTTKKQATSQPNQTPPHEEGDPYFTDLPDNPVVLTVRKPGSKKPVQILALGKPPHRKYFEKV
PEPGKPDRFTLKPINPNGKPGEPSKVYPGGNKPGSRYPVIEYGKPPKAFEVRPDPLHPNDPTKVQFVPVKKVPG
TDGPSFVPAPRTVGKYHTTRKKHHKTTKEPKEGSVIYDDGPYLPKFYPGKKNTKIVAVGKPPKRRYFKIIPKPDKK
FTLKPIDPQNPDDPNPHPEDTKVYPQPDGHPVFEIPGPNKKPEYFKLVPKKDNPKEMDFIPVRPVSPGSFTSKP
RTTRHYKTTKKSKRTTRPTSRVIFEDIPHNTQVPKKRPNQEHPSTIITVGNEPNQKHFEVIPDPSGNPKKFTVVPY
NPYHPNKPCDPKNPKTPQVFEVPKDYPVIQGPDKKFYRLEPDKHNPGQYTFVPVTPLDENHKPSNPHKPDVSF
VPIPRKWHLRTTPGIDDWLDLSTTPSPKTTTDNLGTGYSVVTKPEEISEPTTEVDNAETTVGEEELNKEPTEGPM
TTEATEQFSTEQPTVPSHTETEVAIELTTAEEIILSTASVTQPTVAGKTESTPAELTGKSEIPQSTVSTTEAPTVATH
TPMSVTTEAIQGTTQGAGVSTESETKGQPTGATSGPEEPSKQEPTTSEVSGTGQTASPATSSEAHPRPTAPNV
DDVCFKAYDDCKRSRSSYA*

>**ES_P2** (**hydroxyprolines are highlighted in grey)**
MEMMYTLFFLLFGIVHGQGDGWVLQPDGSYMSYGDGSSGGSYGSTGGSYDGSGGLYGGSSGGSYGELGSG
GLFGGSGGGGFGPGGSYGGFDGGLGGSSGGSQGLPGNGWILQPDGSYLKYEGSGGGGGGGGGGGSGSD
GPPGNGWILQPDGSYMKYDTSGEGNQGSAGSYGGNLPPGFVPPPGYSGNEGGWVLQPDGTYMKTIEQTEP
PTEIRYPVNPDYQPDETLESGKKGSPGYSQVVGFGEPGNMNYFKITPGDAENLYDYSVEPVVSRDNPTHSTDT
EHGKVISRGTKGEPDYQPILQVGPRNHRRYYKMLPDSSKPSGFNFIPQRLTGWLKKQGGDDTSGEVLEPEESE
VTEEANEEIPDEPEIEKKNNDFIICIGRPNRRYLRLKRYSNNPHDYSVEPIGSHDNPDSKPNPNDEEAPEVMTEG
TKGNPNYRSIIAVGPKYNRRYVEVLPNSNDPDKYSFAPVDVQVQPGEQATEPNEEDVEYDDLPENPEYKTIPG
KKKSYEIFSFGPKNRRVYFRIIRRNPNNPNDIEVFPFKDPEHPDSPPSSDDPSAPEVIQQGDPNDPEDGPVIAVG
PKNKKVYFRLGIRKDKVPKFTPVRRGKRIPGKHRLVFTKKTRIWHSRPQKKPLVRKPSAPLKFETSAPVEAATDY
SPDEVDVNPENPVEKRPYQIITFGPPKKRIYIKKTPGPSGNPYDYKLEPIGNPEDPNSKPDPKDPDSPEVIETGEP
GTPSFAPVVAHGPKQNRKYVIIRPNSENPKDPNQMEFSPAEPVPGSDPKKPRFRILRAAVPVAPKDQFTDNTP
KNPNIKTALKPGDKTRYEVISFGPKNRRIYLKKTPGPSNKPDDFKLEPIGNPNDLNSKPDPNDPEAPEVIENGQP
GTPDYSPVVAFGPKTRRKYIIVKPSPKNPKEIQFIPAEPEPGSDPKQRRFKPLTKKVLPGLIRPKPPTHPDSPEIKTV
EGPHPHQVISFGPENKRIHVKKSPGPSGDPNDYKLEPLENPFDPNSKPDPQDKDAPAVIETGKPGTPNFLPIVS
YGPHDNRKYMSFKPIGNTIALSPATLVPGSDPKHPAFNIKTTHPVSPQIPSILKDLHIPTHPLIPPIFNQLPPILNPR
GINLLPGLLPVISNLAELTKPTVQNVEKPNKPSYQVLSFGPLLNRIHIKKTPGPSGKPDDFKLEPIKDPNDPDSKPD
PKDPQTPEIIESGKPGTPDHQQVIAFGPKPQRKYVLVKLNPTDSKKLELSPVEPEPGSDNKQPKFRPVVKPVIPT
LPLPSLDLHNPVINTVEKPNKPHYQVISFGPSTQRIHIKKTPGPSGKPEDYKLEPLGNPDDPDSKPDPKDPNAPEI
VDSGTPGSPDYEPVVAYGPKTARKYVILKPVPLNPKALDLQPAVLEPGSDPKKKKFKVVIQPGISPLLPQILPPFW
PKPSIPYLPKPPPITLPLISHGLQNPIVNNIEEPNKPHMQIISFGPILTRVHIKKILGPTGNPNDYKLEPIGNPDDPD
SKPDSKDLDAPEIIENGKPGTPDYKPVVAYGPKSLRKYVTLQSRPSNPKEIDLEPAELAPGSDAKKPIFKYLPAIPV
FGNKDRNAPEVKTVTDPISKKPVQVISVGPIFKRVHLKKIPGPSGKPNDYKLEPLGNPDDVNSKPDPKDPDSPEI
IENGTPGTPDYAPVVAYGKKIRRKYLKIKSNPLQPNDPTKLLFNRVRLLSPKPGEKTPTFSESLIPKFNPVCYKQYQ
DCNDRS*

**>ES_P3**
MRMKIIILILSLCATTFSSDVTEVADDANPDDLLAQGYVLQPDGSYLKTDETSSTSEQSGTSDQSPDVLKSQGYV
QQPDGTWLKTVEDTETTYDDNEYKNTDYSDGILTVQNDPDVGYFKINKGSSDNENDYSVQAVNSPSDPSPIN
NPGKDDPRVYPQGNQGDPDYHNVIGVGKGKDRTFLKMVPEPSRPSGFKFIPQYISGYRPGDVTNAPEPAPEP
APETSPGSTDDYEEEPNDNPDEPNISDRKGYFIISIGKPNRRFLKLTPSTNNNPHDFSLVPIKSPSEHNSSPDPND
EEAPEVISNENRGSPDYQPVIAVGPPDNRRFTGVVVNPENPKSFQFVVRDQELQPGEAQTEPPEDDIEYEDEP
PNINYKVITVGGKKNEVISAGPKNQRFHFRITHKSKNQGDIEIKAINSADNIDSSPDPDDINAPEIVQQGNPEDP
NNAPIVGCGPRFRRTYHRLIITKGRARFLPLRRVKIRGRHSFIKRNRRWHYRRGIKKIARRVIKRRPLLIRRRKVKY
NKRVVMHRNKLISHNRGGLLIGHKSGHSLGGKGGRLIGGGRAGNGMRHSSRVVIRHHSGSKQNGHGLSLGS
RSEGHHSLSIGGRGSSNGGGGGSHHGSHHG*

**>ES_P4**
MLGKTILILLSFCLFIERSYSFMCRCEKLYRRDMVDGKLVVTEINLCPKTVTEKDCPRGLVRNGCGCCPECGKDL
GQSCSNAMLGPGKCGTGLECVGWEEGNMENTIKAGVCQLKK*

**>ES_P5**
MWNCVRSNNGIMVLLSLLTVKTPITDADSIRALACRALVGLARSEMVRQIISKLPLFTRGELQVLMKEPVLQDK
CLEHVKFCKYASELIEKVTGKPLSSHIESSLAKINKADVVAQTKIVFNEKELLQLIYQHLMNKGYHESALSLEKEAS
LPKNGFTVPGFFGSPPSSSKLARYLFTSVPSASTLSPSIRNHSNTLGGHGSTPIKMNFLTSKNPHNGLNSNKNVK
FKVIRQKSSCGEFQYSPIMKKQNLVKPQLPTVSLDAIITEYLRKQHEHCKNPVVPCPPFSLFVSHHCPDPMFRNS
APNNITARILRRSAFPKHGGIDGARLNRKHIYSRFRPVRSYRGVEEEGCFCCCAFSHDDENLYLGTYTGEIKQYNI
QTRTEEASYNCHTSALTLLEPSQDGKLMLTSASWGRSLSGLWAFQNNSLELMYSFENDHFVEFSKLSQDRIIGT
KEETAHIYDVSTGQLVRTLYNADLANNYTKNRATFSPFDDLALSDGVLWDVQAANPIHKFDKFNPHISGVFHP
MGLEIIINSEIWDLRTFHLLHTVPALDQCQIVFNSAGDVMYGANHQLDDDNDLSEDSVKSPFGSSFRTFDATD
YSNIATIDIKRNIFDLSTDKSDCLLAIIENQGARDGLVEESICRLYEVGRTKDQDDEQVGGAAGGAGE*

**>ES_P6**
MGAKYKPDWEDSCTHLICAFANTPKHRQVQKLGGQVVRKEWIVECYKKKRKLPWKMFRLDGDSEEEEDDED
NDQTYEEGDDEEDEDEEEIRLEKKYSKKHVPKPGPSNLKKTSPKKTTLTFTEKKPSPKKLKLYDEEENEENSKKYS
SEEDSGDDTEDEIRKVREEENKKPNIQHTKSDNEYEGSTDEDAELEKKNVKSLSKTIVENGNSSSTKESIFPLPNL
PSFFKEKSFFFFGNFDDATRKLLKRYIIAFGGKLENSMDQKVSFVITASDWDEQFEQALNENDSLTFVRPQWIF
KCKDKGKFVPYQPYVVVPKESSD*

**>ES_P7 Phosphorylated sites highlighted in grey**
MKTIIWFFISICVADSAIRVSHQYNFKEKPKNQNSRNSIKRTSGGYGAEMDNTGVLINKYPDTPPPFYQMLPDD
FMNPPDYNNNNNNNGGGGINPPTDYSSGSFDPMPPDDFMNPPDYNNNNNNNGGGINPPTDYSSGSFDP
MPPDDFMNPPDYYNNNGDINPSTDYNSDSFDPISVPDYNNNNGDNNNNQENYENNYNQNLGNNEQIDEE
MPPDYVSSDSGYLPPGYIAPPGISRDGWHLLPDGNYVKYIVETEAPIAYEVPISPVDNPEILPVGEAGSPGYHDII
GIGEPNIRSYFKRTPGPSGKPDDYTLEPIVSPRKPIPVNPKLRNRPRVIYPEGRNNPPTLMIGPEGRRKYFKMVP
NDDKPGGYDFIPRIKKGSKIPEPSHIITQKPQISETTEETELTTTELLEPPTEEVTESPDNYDEEPEFDDLPENPQFE
THRPGTNHPFTIISLGPKNHRQHFKKIPRSRNPNDYDLEPYDPNTPNHKVNPRNPYAPKVLNNGIKGTPQFSPV
IKTGQPHFPKFYRLVPHPNNPNQMGFVPVTKNILKPGNKKPTFSDLPRQFKPPRRRRPEEVRDDEPDDGEPEF
SDLPEEPHFKTVKQPKRHPYTLISVGKPPHKNTFKKTPGKSGNPNDYTIEPFDSDTPNNEPDQDNPDPPKVHP
HGEKGKPNYSPVIEYGPHKHRKFFRLVPHPTHPNHFGFIPVKKVNTKKGPHFQDLPRKYQPHRIHSPTTHKPHE
GSKVFDDLPDDPSIITRHLPGRRHPTQIISVGKPPHRRHFEKKPGPSGDPDDYVIEPFDPKNPNKKPNPHNPKSP
KVYHGKHPVIKTGSPQKPKYFRVEPDPNHPHDPNRVKFTPVKPLKKGKNPTFVEIPRRFHIYHTTPKSSEVTTTE
PEDDEPVYDYQPERVKIKNIHPPHSKYPYQIISIGKPPNRQHFLKKPGPSKKPYDYTIEPYDPKTKDHKPHSKNPP

KVYPAKGIPGNPDYQPPVIRVGPKNKPKYYKLKPDPKHPHDPRKIKFIPVIPVPHRPGKFQNLPYHYYHRTTTKE
PDESDIIFDDLPENPEVKELKRPHQKIPSQIISVGKPPYRKHFEKKPNPSGKPDDFEIYPFDPMHPNKKPKPSDDV
FPATGKPGTSKYQPPVIKTGSPQKPKYFRIIPNPDYPNDPKKIIFIPVKPLKRPDSHKLNFIPLKRHYHIYHTTTKKH
PKPPRGQTPEPYDGEPIFVDQPENPKVFQVKDPKHKRPVQIIAVGKKPHRQYFKKIPGPSHKPDDYRILPFDPR
KPNGKPDPKDKSSPKVYPGPPPVIEYGPKHNRKTFKLVPDPKHPNDPTKINFIPVKKISNKPYDNKPHFQNLPRL
TRLYHRTTKHKHPKTTAEPQEGDIVFTDEPYKPKIYTVKEPGRKHPVQIIAVGKPPNRKYFKKVPKPNDPNDFILY
PYDPQNPHENPNPEDDKTPKVYPGDKNPRKPKHRFPVIETGTPQNPNFYEVVPDPKHPGKVNFVPVIPVKNIE
KPEKPLYFIKPTRKFYRYTKSKKTTTPEPKEGEPQFIDVPDTKVEQKVVPGYKKPVQIVEVGERPYTKKFVKIPGPS
KHPDDYILKPFDRKHPEKNPKTPKVYGPFGKPNTPDYHHPVIEFQPNKFFEVKPNPKYPHDPKKIILVPVKPTKP
KDKNPSFVELPRKWYCYHTTFEPEKEIIEGTTNPEKLTTIQEATTEELITVNDGTIVTITEAELTEELTDNPEKKESE
VPEKTEEPEFNSTPKPNQSEEVEEPEEPGYPKPQKPSGTGEAEEPERSEESEEPGQKPRHPKPHRPGSGGEPEM
PGAENEGPEEPEKPREPGTGEQEKPGEPETEEPEYKPRHPKSQRPGEENEEPKEPEKPREPGTGEPEGHEELEK
PGEPERPGNKPRHPKPQRPGGENEEPEEPEKPREPEGSGHKPQRPGGEPGEPEEPGQPEEPGHKPRHPKPQK
*
